# Home range spillover in habitats with impassable boundaries: Causes, biases, and corrections using autocorrelated kernel density estimation

**DOI:** 10.1101/2024.11.20.624379

**Authors:** Jack P.W. Hollins, Christen H. Fleming, Justin M. Calabrese, Les N. Harris, Jean Sebastien Moore, Brendan K. Malley, Michael J. Noonan, William F. Fagan, Jesse M. Alston, Nigel E. Hussey

## Abstract

1. An animal’s home range plays a fundamental role in determining its resource use and overlap with conspecifics, competitors and predators, and is therefore a common focus of movement ecology studies. Autocorrelated kernel density estimation addresses many of the shortcomings of traditional home range estimators when animal tracking data is autocorrelated, but other challenges in home range estimation remain.
2. One such issue is known as ‘spillover bias’, in which home range estimates do not respect impassable movement boundaries (e.g., shorelines, fences), and occurs in all forms of kernel density estimation. While several approaches to addressing spillover bias are used when estimating home ranges, these approaches introduce bias throughout the remaining home range area, depending on the amount of spillover removed, or are otherwise inaccessible to most ecologists. Here, we introduce local corrections to home range kernels to mitigate spillover bias in (autocorrelated) kernel density estimation in the continuous time movement model (ctmm) package, and demonstrate their performance using simulations with known home range extents and distributions, and a real world case study.
3. Simulation results showed that local corrections minimised bias in bounded home range area estimates, and resulted in more accurate distributions when compared to commonly used *post-hoc* corrections, particularly at small-intermediate sample sizes.
4. Comparison of the impacts of local vs post-hoc corrections to bounded home ranges estimated from lake trout (*Salvelinus namaycush*) demonstrated that local corrections constrained bias within the remaining home range area, resulting in proportionally smaller home range areas compared to when post-hoc corrections are used.

## 1. Introduction

The estimation of an animal’s home range is often a key objective of electronic tracking studies due to the fundamental role of the home range in resource selection (Slaght et al. 2013) and interactions with conspecifics, competitors, and predators (Martinez-Garcia et al. 2020; O’Brien et al. 2020). There are many analytical approaches to home range estimation, each with associated assumptions, caveats and limitations (Signer and Fieberg, 2021). These can be broadly grouped into geometric and probabilistic methods. Geometric estimators, such as the minimum convex polygon (Burgman and Fox, 2003) and local convex hull family of estimators (Getz et al. 2007; Lyons et al. 2013) generate home range estimates from polygons constructed from groups of recorded animal positions. In contrast, probabilistic home range estimators assume and target an underlying probabilistic model when constructing a home range and can be further distinguished as either parametric (e.g., mechanistic home-range analysis [Moorcroft et al. 2008] or Autocorrelated Gaussian Density Estimation [Fleming et al. 2015]) or non-parametric (e.g., kernel density estimation [KDE]).

In (A)KDE, individual density kernels are placed on each animal relocation in the dataset (constituent kernel) with the total home range provided by the sum of all constituent kernels (Silverman et al. 1998; Fleming et al. 2015; Fleming and Calabrese, 2017). The width of these constituent kernels is referred to as the bandwidth, and the method used to optimise the bandwidth controls the final size and shape of the resulting home range estimate. Smaller bandwidths yield multimodal range estimates with peaks tightly associated with clusters of recorded locations, while larger bandwidths smooth over these peaks and yield a more spread-out distribution. The method used to optimise the bandwidth distinguishes the different methods of (A)KDE-based home range estimation, with each optimization approach making different assumptions and approximations of the underlying data (e.g., Silverman, 1998; Worton, 1989; Gitzen and Millspaugh 2003). Of these approaches, currently only the autocorrelated Gaussian reference function employed in autocorrelated kernel density estimation (AKDE) accounts for autocorrelation in animal tracking data during bandwidth optimization (Fleming et al. 2015; Calabrese et al. 2016; Noonan et al. 2019). (A)KDE is thus one of the most commonly employed means of home range estimation, including situations where animal movement may be constrained by impassable boundaries (e.g. Mate et al. 2016, Wells et al. 2017). As constituent kernels spread from each individual location, the home range estimate will extend beyond the distribution of animal relocations by a degree proportional to the bandwidth. Consequently, where animals move near impassable barriers, such as cliff faces, shorelines or roads, home range estimates may extend into unavailable and unused habitat (‘spillover bias’; Knight et al. 2009). Spillover bias can therefore lead to overestimates of home ranges calculated via (A)KDE where animal movement is constrained by the presence of impassable boundaries, with the bias introduced being proportional to the size of the bandwidth and the amount of time a tracked animal spent near that boundary (Noonan et al. 2019).

Several approaches have been developed to improve home range estimates influenced by impassable boundaries, including incorporation of ecological processes (resource selection, competition, etc) to link home range estimates directly to underlying environmental conditions (Moorcroft et al. 2008, Slaght et al.2013, Matthiopoulos 2003; 2003a), or post-hoc corrections to more traditional home range estimators (e.g., Fury and Harrison, 2008). However, these approaches are often computationally expensive (Péron et al. 2019), only work under specific conditions, or create tradeoffs with other aspects of home range estimation (Noonan et al. 2019). In this paper, we provide an overview of different approaches used to accommodate the presence of impassable boundaries in (A)KDE-based home range estimation and associated spillover bias. We also demonstrate a novel extension of the existing AKDE framework that accounts for impassable boundaries (e.g., coasts, fences, roads) to movement, which we term locally corrected AKDE. The overview discusses the strengths and weaknesses of corrective approaches (i.e. those which remove spillover as it is generated), and those used to prevent spillover occurring when estimating home ranges via (A)KDE. We focus on home range estimation via (A)KDE as these approaches are widespread in ecology, while AKDE is the only home range estimator designed for use with autocorrelated tracking data. For researchers interested in the application of non-KDE-based home range estimation, we have briefly reviewed the application of geometric, mechanistic, and lattice-based home range estimators and their capacity to account for impassable boundaries in Appendix S1. We strongly recommend consulting this material when researchers are considering the use of non-KDE based home range estimation and the impact of impassable boundaries is a concern. We then quantify the performance of locally corrected AKDE using simulated animal trajectories with known home range extents and distributions constrained by an impassable boundary, and compare it to performance of commonly used *post-hoc corrections*. Finally, we illustrate the performance of locally corrected AKDE and the importance of considering spillover bias in home range estimation using fine scale acoustic tracking data from lake trout (*Salvelinus namaycush*) collected from two high-Arctic lakes with contrasting shoreline complexities.

### 1.1 Correcting spillover bias in KDE-based home range estimation

In (A)KDE, a home range is estimated by summing constituent kernels placed on each individual location and normalising the total density contained by all kernels such that it integrates to 1. The coverage areas of the final probability density function (PDF) then provide the home range estimate’s area. Integration of the PDF allows calculation of the corresponding cumulative distribution function (CDF). While the PDF describes the probability density of a tracked animal’s range distribution (Alston et al. 2022), with each cell of the distribution providing an estimate of ‘intensity of use’ (i.e., all values are >0, and integrate to ‘1’ or ‘100% use’), the CDF defines the relationship between a given density isopleth (i.e., that pertaining to 50% or 95% density) and the area that this isopleth encompasses. Where animal movement is constrained by a boundary, corrections may be applied to constituent kernels (local corrections), or to the PDF or CDF (post-hoc corrections), to remove spillover bias.

#### 1.1.1 Local Corrections

Local corrections are applied to constituent kernels before PDF integration. One example of local correction was demonstrated by Benhamou and Cornelis (2010) using animal relocations that fall close to a movement boundary to generate reflected ‘mirror images’ of these positions onto the opposite, unavailable boundary side. By then placing negative density kernels upon those positions, the outermost contour of the final PDF will fall equidistant between the ‘true’ location on the available side of the boundary and the corresponding reflected point on the opposite side. The resulting home range estimate therefore conforms to the boundary, while also localising the impacts of this correction to regions close to the boundary itself (Fig. 1). However, this approach is only viable where boundaries consist of simple, straight-line segments that are much longer than the home range estimate’s bandwidth. This approximation will therefore not hold where boundaries are complex or when large optimal bandwidths are required. An alternative local correction to kernels was developed by Guo et al. (2019), who leveraged underlying environmental information to control how kernels spread from individual locations, so that their shape adhered to the presence of impassable boundaries. In this approach, bandwidth selection is accompanied by optimization of a topographic influence parameter which dictates the shape and spread of each constituent kernel according to its underlying topography, preventing extrapolation across impassable boundaries.

**Figure 1:**
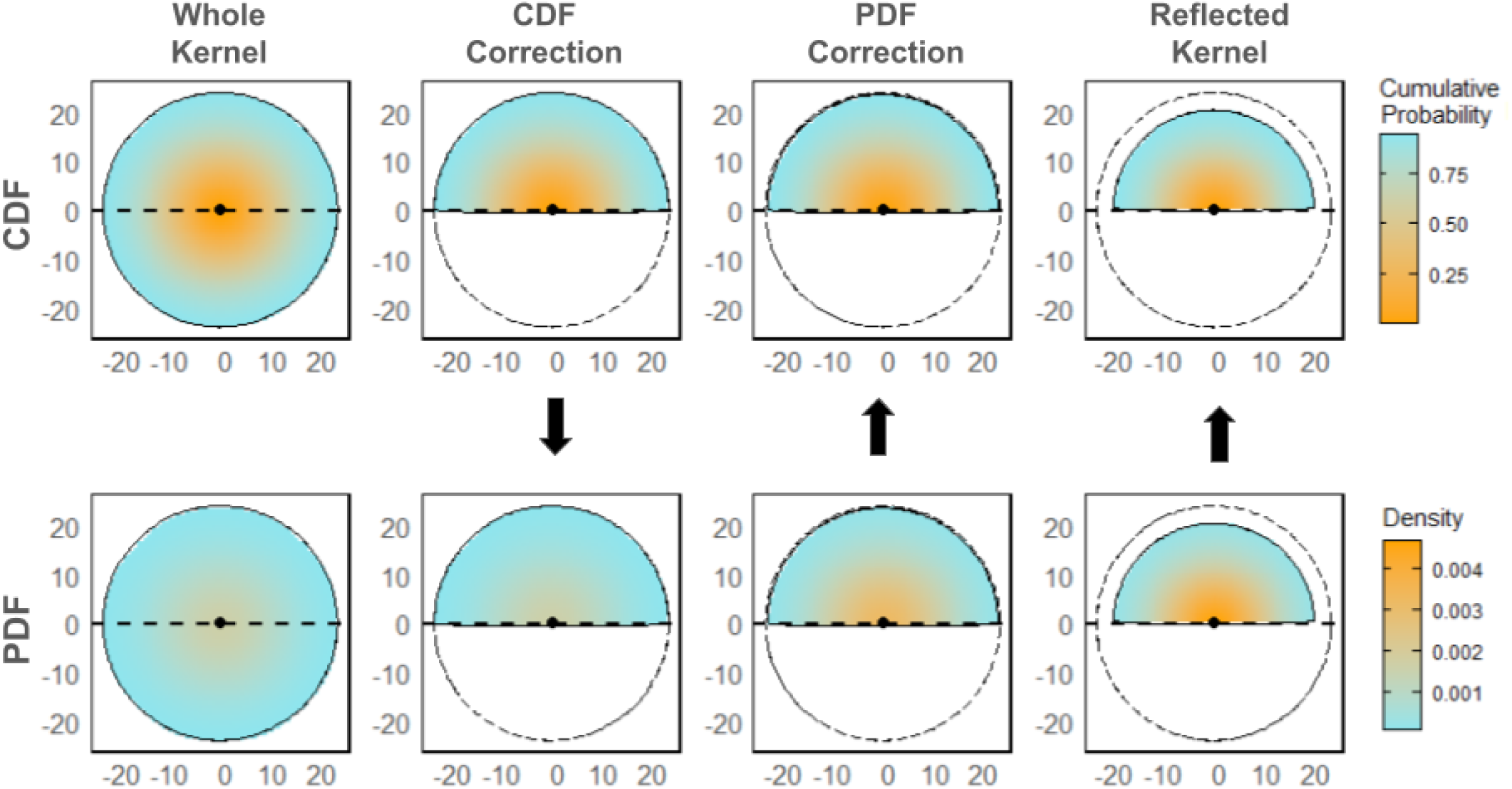
Impact of post hoc correction of the cumulative distribution function (CDF) and probability density function (PDF), and the local reflected kernel correction on the CDF (upper row) and PDF (lower row) of an individual home range kernel. Where correction was applied to the CDF, the corresponding PDF was calculated, and vice versa (black arrows). Animal position is indicated by the black dot, and the area corresponding to the 95%, untruncated kernel (far left column) indicated by the circular dashed line. Both post-hoc PDF and reflected kernel corrections redistribute excluded cell probability densities on the available boundary side, while the reflected kernel is also contracted compared to the uncorrected kernel. In the case of post-hoc CDF correction, probability densities show minimal change from the uncorrected kernel, but will lead to positive bias in area when kernels are integrated into a home range estimate.

#### 1.1.2 Post-hoc corrections

*Post-hoc* corrections to the home range extent are the most frequently used methods to remove spillover (Knight et al. 2009; Lapointe et al.2013; Hawley et al. 2016; Mate et al. 2016; Wells et al. 2017), and are often also referred to as ‘masking’ or ‘clipping’. Post-hoc corrections truncate either the CDF or PDF at an impassable boundary, each removing inaccessible areas but differing in how spilled-over home range is redistributed within the available area. In CDF correction, the removed probability mass of the spillover is redistributed to the isopleth from which it was removed on the available boundary side. In contrast, when correcting the PDF, the removed mass of the range estimate is redistributed throughout the entire remaining home range extent, proportionally increasing the probability mass contained within each remaining cell (Fig. 1). While post-hoc corrections provide a more biologically relevant range estimate compared to the uncorrected home range impacted by an impassable boundary, these corrections are not localised, introducing bias throughout the resulting home range estimate by a degree proportional to the amount of spillover removed (referred to as ‘redistribution bias’ hereafter). Post-hoc corrections to home ranges are compatible with both simple and complex boundary shapes, are straightforward to conduct in analytical software such as R, and are also supported in other geospatial software (e.g. ArcGIS Pro 3.3.1, ESRI, 2024).

### 1.2 Preventing spillover bias in (A)KDE-based home range estimation

While local and post-hoc corrections to home range extents account for spillover after kernels have been placed, the presence of an impassable boundary can also be incorporated earlier in the home range estimation process to prevent spillover from occurring in the first place. This approach was demonstrated by Péron (2019), using an extension to AKDE that parameterized animal-environment interactions using a movement model with a weight function describing selection of specific habitat types and movement barrier permeability. These animal-environment interactions were then used to condition bandwidth optimisation upon estimates of habitat preference or the tendency of tracked animals to cross movement barriers, yielding optimal bandwidths which respect impassable boundaries. Inability to cross a movement barrier was controlled using a permeation term, implying straight line movements along boundaries. Péron also included these environmental interactions at the constituent kernel placement stage, where kernels are truncated at the impassable boundary. Although this provided a robust method to prevent spillover bias, Péron (2019) found this method of bandwidth optimization computationally prohibitive, and so also demonstrated a simplified version, which only leveraged the environmental weights quantified at the model fitting stage to shape the final range distribution, termed E-AKDE and SE-AKDE for the full and simplified versions, respectively. Péron (2019) also noted that the straight-line movements implied by the permeation term would not be suitable where boundaries are more convoluted.

### 1.3 Preventing spillover bias using transformation-based home range estimation

Spillover bias results from the continuous kernel functions used within (A)KDE, which effectively assume that the area encompassed by a constituent kernel is equally accessible to and used by the tracked individual. Home range estimates based on (A)KDE therefore ‘smooth over’ areas which are close together in Euclidean distance, irrespective of their habitat suitability or availability. Spillover bias can therefore also be addressed by transforming Cartesian animal locations into locations described by environmental covariates such that animal locations occurring in more ecologically similar locations are closer together in the resulting distribution. The home range is then estimated via KDE on these non-Euclidean locations, smoothing across environmentally similar locations and preventing extrapolation into areas unused by the animal. After assigning probability densities to these locations, they are then back-transformed into their Cartesian locations along with their corresponding density estimates. Resulting home range estimates therefore adhere to the presence of any impassable barriers, assuming the barriers are adequately described by the environmental covariates chosen for transformation.

Transformation-based estimation was used by Tarjan and Tinker (2016) to estimate coastal home ranges of sea otters (*Enhydra lutris*) in California, transforming latitude and longitude positions into locations defined by their distance from land, and their position along the 10m coastal isobath. Distances from land were log transformed, assigning terrestrial positions a value of log(0), which is infinitely far away from any sampled location, excluding these areas from the home range. A more linear transformation was used by Péron *et al*. (2017) when investigating the probability of collision hazard between wind turbines and three species of soaring raptor, the Andean condor (*Vultur gryphus*), griffon vulture (*Gyps fulvus*) and golden eagle (*Aquila chrysaetos*) using three-dimensional home range estimates generated from longitude, latitude and flight height, with the ground representing a movement boundary. To implement this constraint within their analytical framework, Péron *et al*. (2017) used a custom-designed link function that was linear across flight range, preserving autocorrelation in velocities, but diverging towards -∞ as flight heights approached 0. This more linear link function was critical within the stochastic process used by Péron et al. (2017), as the log link function employed by Tarjan and Tinker (2016) skewed speed estimates of tracked raptors depending on their position relative to the boundary, such that the movement model became unrealistic, hampering its capacity to account for autocorrelation in the data.

## 2. Methods

Existing approaches for addressing spillover in home range analyses are of limited utility. For example, only a subset of the described approaches to addressing spillover are mathematically accessible, whereas others introduce further bias into home range areas (e.g., post-hoc corrections), or require specific, strong conditions to be effective (e.g., mirrored local corrections). To provide an accessible alternative that avoids these operational constraints, we developed a method to remove home range spillover within the AKDE framework (available in the `ctmm` R package, Calabrese et al. 2016). This method retains the ability of post-hoc corrections to work with complex boundaries, while constraining the redistribution of spillover to the area adjacent to the boundary.

### 2.1 Locally Corrected (A)KDE

When using locally corrected (A)KDE, bandwidth optimisation and placement of constituent kernels occurs as described in Fleming et al. (2018) and Noonan et al. (2019), but the PDF of constituent kernels is truncated at the movement boundary where spillover occurs. This removed probability mass is then redistributed within the constituent kernel it was removed from before PDF integration. Within the `ctmm` package, local kernel correction is supported within the akde() function call by supplying a SpatialPolygonsDataFrame object corresponding to the shape of the boundary to the ‘SP’ argument, and specifying whether the animal locations are expected to be in or outside of the polygon (SP.in = ‘TRUE’ or ‘FALSE’, respectively). The `akde()` function then rasterises the polygon using a resolution corresponding to the grid on which the home range is calculated, and an extent corresponding to the dimensions of the uncorrected home range. The cells of this raster are then assigned a binary value depending on whether or not they are inside the boundary, which is used to identify portions of the constituent kernels which fall on the inaccessible boundary side. The cells of the constituent kernels on the inaccessible boundary side are then removed, and their contained probability mass redistributed throughout their corresponding constituent kernel. Locally corrected AKDE can therefore remove spillover bias for home ranges with more complex boundary shapes compared to simpler shapes required by the mirrored local corrections method introduced by Benhamou and Cornelis (2010). However, locally corrected AKDE will still leave small amounts of spillover where boundary segments intersect the cells of the home range estimate. Where this is a concern, users may specify a finer grid resolution within the AKDE call, although this can increase computation time.

### 2.2 Demonstrations of Locally Corrected (A)KDE with CTMM

#### 2.2.1 Trajectory simulations within a hard boundary

To assess performance of local kernel corrections vs post-hoc PDF/CDF corrections, we applied AKDE to simulated animal movement data with an impassable movement boundary. These simulations provide simple but realistic movement scenarios where true home range sizes are precisely known and therefore can be used to check the quality of the home range estimates. Simulations were carried out in R (Ver 4.4.0). using the `ctmm` package (Ver. 1.2.0) following the approach outlined by Noonan *et al*. (2019), with corresponding code and further details of the simulation included in Appendix S2 of this paper. In all simulations the boundary was identical and enforced along *y*=0.

Two simulation approaches were used to investigate overall performance of local kernel corrections. The first set of simulations consisted of a total of 10,000 trajectories with a fixed tracking duration of 32 days with one home range crossing per day, and eight positions recorded per day. The second set of simulations varied the tracking duration (2, 4, 8, 16, 32, 64, 128, 256, 512, 1024, 2048, and 4096 days), with 400 trajectories simulated for each tracking duration. For each simulated trajectory, a corresponding home range was estimated using AKDE, and spillover removed using post-hoc PDF and CDF corrections, and the newly introduced local corrections (Fig. 2). The true area of each bounded home range was then calculated using the exact area relations described in supplement S3 of Noonan et al. 2019, to which the areas of the different corrected home range estimates were then compared. This provided a means to compare the bias in home range estimates resulting from each correction approach. Bias was determined by calculating the proportional difference between the true and estimated 95% home range sizes, and expressed as a percentage (% bias). For all statistical comparisons, and throughout the text, percent bias is expressed as an absolute value, but signed values were retained for plotting purposes corresponding to overestimates and underestimates, respectively.

**Figure 2:**
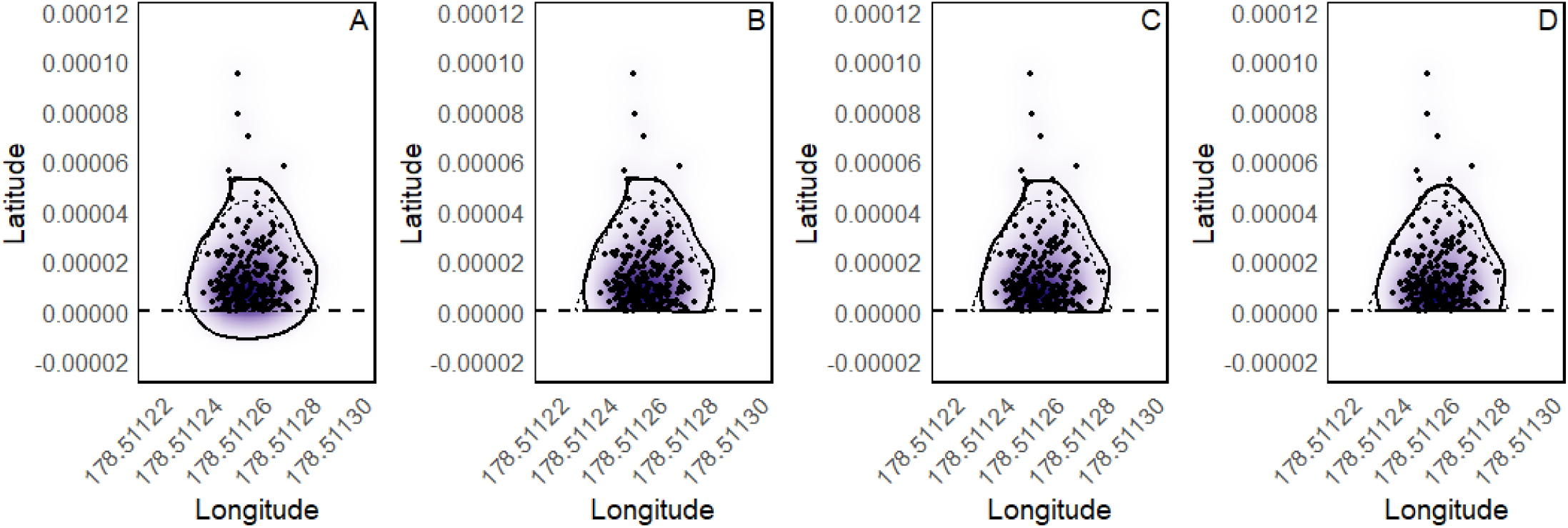
Comparison of post-hoc and local correction of AKDE home range estimates, overlaid on the true home range (dashed line). An impassable barrier exists and Latitude = 0. A) shows the home range estimate with no boundary enforcement, while B, C, and D show the impacts of CDF, PDF and local correction, respectively. In this example, the three correction methods (CDF, PDF, and local) yielded home range estimates that exceeded the true values by 19.4%, 16.6%, and 12.7% respectively.

In addition to the percent bias of each correction type, the similarity of corrected home ranges to the true density distributions was assessed using Bhattacharyya distance (BD, Bhattacharyya 1943; Winner et al. 2018) which compares the similarity of two probability density functions. Bhattacharyya distances of zero indicate that there is no difference between the true and estimated range distributions and increase as the two distributions diverge. The true density distribution of each simulated home range was constructed using the grid coordinates of the corresponding AKDE and true cell densities determined using the normal-χ^2^ distribution used to simulate bounded trajectories (Appendix S2, Noonan et al. 2019).

Differences in percent bias and BD between the different home range corrections were examined via ANOVA. In addition to comparisons to known home range extents and distributions, simulated home ranges across different tracking durations were also compared to uncorrected AKDE home range estimates generated from the same simulated data. Differences in percent bias and BD between the different methods for correction across sample sizes were investigated using a generalised additive model (GAM, `mgcv::gam,` Ver.1.9-1, Wood, 2011) to model both percent bias and BD, using correction type and tracking duration as predictors. Post-hoc analysis of GAM results was performed via `emmeans::emmeans (`Ver.1.10.4, Lenth, 2024), and GAM fit assessed via `mgcv::check.gam`. For simulations with fixed tracking durations, we also calculated the proportional difference in estimated 95% home range size when CDF or local corrections were used, and its relationship with the corresponding trajectory’s mean distance from the boundary. This relationship was used to investigate how the impact of different boundary corrections on home range estimates depends on the movement behaviour of tracked animals using a linear model via `base::lm.`

#### 2.2.2 Case Study

To demonstrate the utility of local corrections to kernels in a real-world ecological scenario, we used acoustic telemetry data from lake trout tracked using a Vemco Positioning System (VPS, Innovasea, Halifax, NS) over an entire year. The study took place at two High Arctic lakes (Inukhaktok and Nakyulik) on southern Victoria Island that displayed contrasting shoreline complexities. A total of 20 trout were tagged (n=8 in Inukhaktok, n=12 in Nakyulik), and their home range estimates calculated via AKDE. One Inukhaktok trout spent the entire tracking period sufficiently far from shore to exhibit no spillover in its home range, and so was excluded from subsequent comparisons. Home range estimates then had spillover removed via post hoc CDF, PDF and local corrections, and the resulting impact on home range size compared by calculating the proportional reduction in uncorrected home range size caused by each correction. The proportional reduction in home range size caused by each correction approach was compared using a general linear mixed model, using correction type and fish ID as fixed and random effects, respectively (lme4::lme, Ver3.1-166, Pinheiro et al.2024), with pairwise comparisons performed via `emmeans::emmeans (`Ver.1.10.4, Lenth, 2024). The relationship between home range overestimation when using CDF correction as opposed to local corrections, and the distance fish spent from shore was also investigated using a random subset (100 randomly selected with replacement) of monthly home range estimates from each lake. Monthly home range estimates were used to allow seasonal inshore/offshore movements to contribute to the magnitude of home range spillover, and provide a wider range of distances from shore to be investigated.

## 3. Results

### 3.1 Simulation Results

#### 3.1.1 Trajectory simulations within a hard boundary: Fixed tracking duration

Correcting for the impassable boundary present in the simulated data via local corrections to kernels resulted in less biassed home range estimates (6.53±6.79%,; F_(2,2997)_=2525, p<0.001, Fig. 3, Table 1) compared to both post-hoc PDF (p<0.001) and CDF corrections (p<0.001), which had mean biases of 10.6±6.63% and 13.36 ±7.07%, respectively (Table 1). A similar pattern was evident when comparing similarity between true and estimated home range density distributions via BD (F(_2,29996_)=4204,p<0.001), with PDFs from locally corrected AKDE more closely representing the true density distribution than both PDF-(p<0.001) and CDF-corrected (p<0.001) AKDE. Mean BD between locally corrected AKDE and the true distribution was 0.13 (±0.07), while PDF- and CDF-corrected AKDE estimates showed BD distances of 0.15 (±0.08) and 0.23 (±0.09), respectively. Overestimation bias of CDF-corrected home ranges was significantly related to the mean distance of the simulated trajectory points from the movement boundary, with the discrepancy between CDF and local corrections increasing with decreasing distance from the boundary (t=-15.04, p=<0.001, Appendix S3).

**Figure 3:**
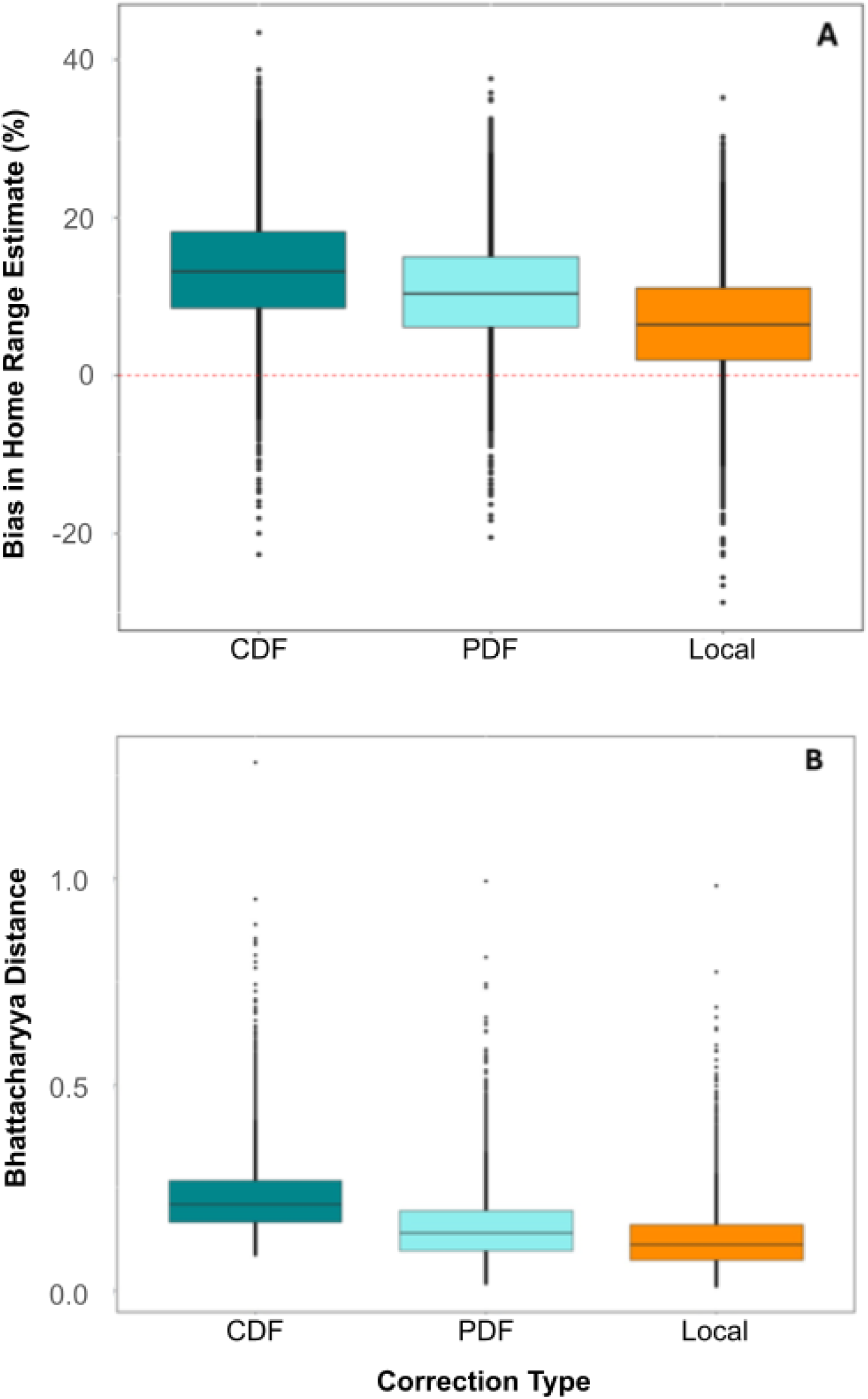
Comparison of the bias of 95% home range estimates calculated for the simulated bounded movement data (n=10000 trajectories) with the movement boundary enforced via post-hoc CDF, post-hoc PDF and Local corrections, in terms of both area (A) and distribution (B). The dashed red line corresponds to 0% bias in panel (A). Local corrections to home range kernels minimise bias in area estimates and yield cell densities which most closely represent the true distribution.

**Table 1:** Summaries of the accuracy (%) and similarity between calculated and true home range distributions (BD) resulting from different methods of enforcement of a hard movement boundary on AKDEs calculated from simulated data.

| Correction Type | Mean Accuracy (% of true HR) | Accuracy SD | Mean BD | BD SD |
| --- | --- | --- | --- | --- |
| *Post-hoc CDF* | +13.36 | ±7.07 | 0.23 | ±0.09 |
| *Post-hoc PDF* | +10.6 | ±6.63 | 0.15 | ±0.08 |
| *Local* | +6.53 | ±6.79 | 0.13 | ±0.07 |

#### 3.1.2 Trajectory simulations within a hard boundary: Variable tracking duration

Both GAMs investigating the performance of the different home range corrections across sample sizes indicated significant effects of tracking duration (percent bias:F=9086, p<0.001; BD:F=509.9,p<0.001), and correction type (Local vs CDF t=33.606, p<0.001; Local vs PDF t=16.899,p=<0.001 and Local vs CDF t=10.453, p<0.001; Local vs PDF t=10.650,p<0.001 for % bias and BD respectively). Similar to the simulations with fixed tracking durations, local kernel corrections to home ranges estimated from variable tracking durations showed the lowest percent bias compared to CDF and PDF corrections, although differences were less pronounced than observed in simulations with fixed durations (mean bias of 17.13±14.33%, 17.79±15.5% and 20.1±18.3% for local, PDF and CDF corrections, respectively (S3)). The positive bias in locally corrected, PDF- and CDF-corrected home ranges each initially decreased with tracking duration, before estimates converged to an underestimate, resulting in negative bias at high sample sizes (Fig. 4A). Locally corrected home ranges reached their minimum percent bias with less data than PDF- and CDF-corrected home ranges, each intersecting 0% bias at approximately 59.37, 71.67 and 100.35 days, respectively (Fig. 4A). The BD between the true distribution and estimated home range was consistently the lowest for locally corrected home range estimates compared to both CDF and PDF corrections across all tracking durations (Fig. 4B). Similarity between corrected and uncorrected home ranges increased with increasing sample size with respect to both their respective areas, and density distributions (Figs. 4C, 4D, respectively).

**Figure 4:**
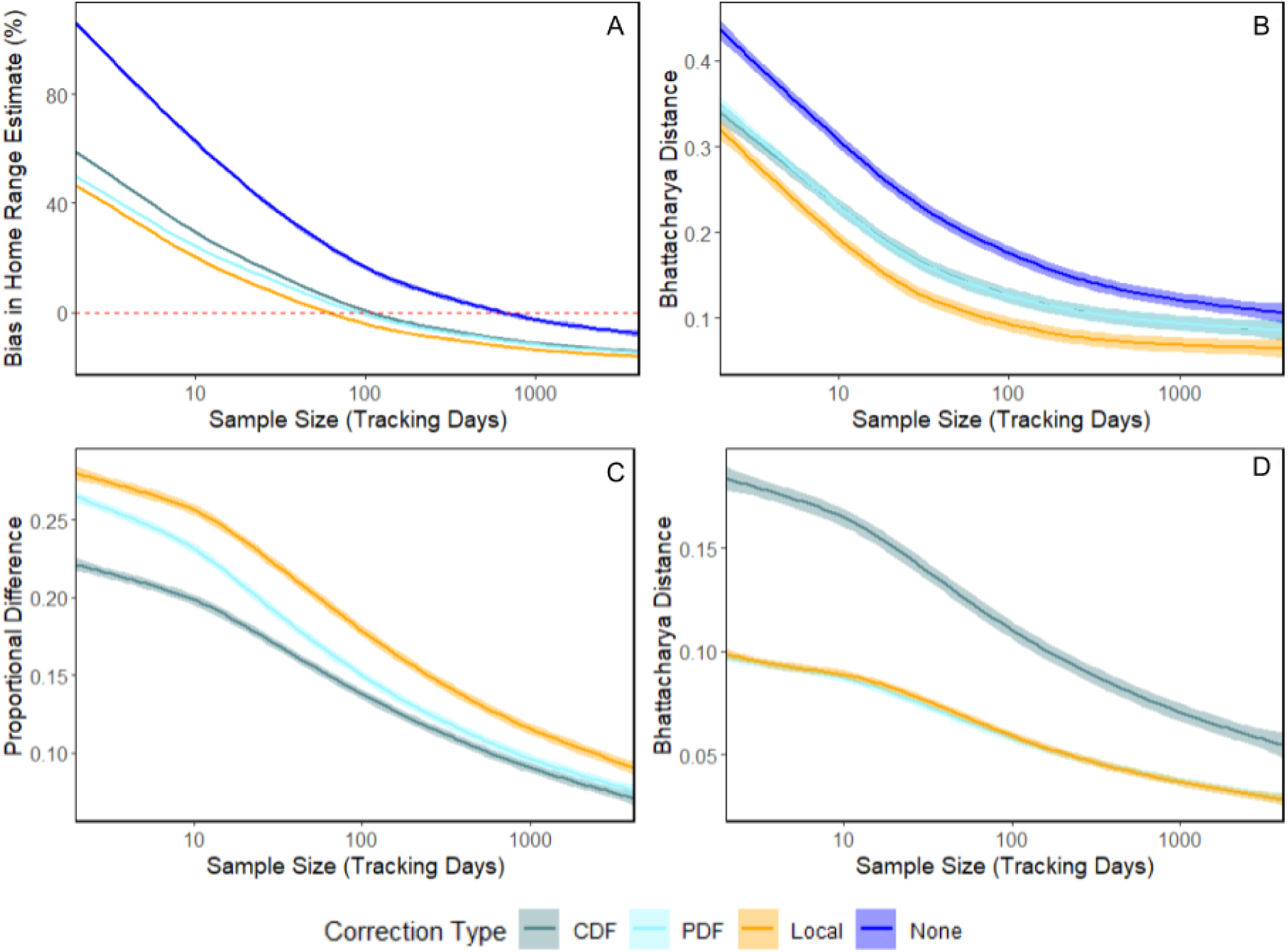
Comparison of the impact of different boundary-corrections on simulated HR estimates provided via AKDE across tracking durations. Panels A and B show bias in area estimate and the Bhattacharyya distance as determined via comparison of simulated home ranges to true values, respectively. Local corrections to kernels provided the least bias in estimated 95% home range areas at short to intermediate sample sizes, but underestimated 95% home ranges at progressively longer tracking durations. Local corrections to kernels resulted in the lowest Bhattacharyya distances, compared to other correction approaches, at all tracking durations. Panels C and D compare the corrected to uncorrected home range estimates across sample sizes. in terms of the proportional difference in 95% home range area, and Bhattacharyya distance, respectively. Proportional differences and Bhattacharyya distances between corrected and rected home range estimates decline with increasing sample size as the amount of spillover decreases, reducing the act of different correction approaches. In all cases, curves were generated using a general additive model (GAM), with shaded areas corresponding to 95% confidence intervals.

### 3.2 Real World Case Study

Results of the GLMMs comparing spillover corrections indicated significant differences in the impact of the CDF, PDF and local corrections in both lakes, with local corrections resulting in the greatest proportional reduction in home range size compared to both CDF (p<0.001) and PDF (p<0.001) corrections (S3, Fig. 5). CDF, PDF and local correction reduced the uncorrected home range by an average of 25.3 ± 12, 30 ± 13 and 35.1 ±14% (mean ± SD), respectively. This greater proportional reduction of local vs either of the post-hoc corrections resulted in local corrections providing the smallest home range estimate in all cases, although in two instances PDF and locally corrected estimates were within 1% of one another. Similarly to the fixed-duration simulation, overestimation bias of CDF-corrected trout home increased with the amount of time trout spent close to shore in both Nakyulik (t=-3.958,p=<0.001) and Inukhaktok(t=-2.346, p=0.02, Fig. 6, S3). Overestimation bias caused by CDF correction has further implications for the proportion of animal relocations contained within the home range estimate, potentially resulting in more than the target of 95% of recorded animal positions contained within a home range estimate (Noonan et al. 2019; SIgner and Frieberg 2021, fig.7,8), particularly where the observed number of home range crossings is low. This impact of redistribution bias is discussed in more detail in Appendix S4.

**Figure 5:**
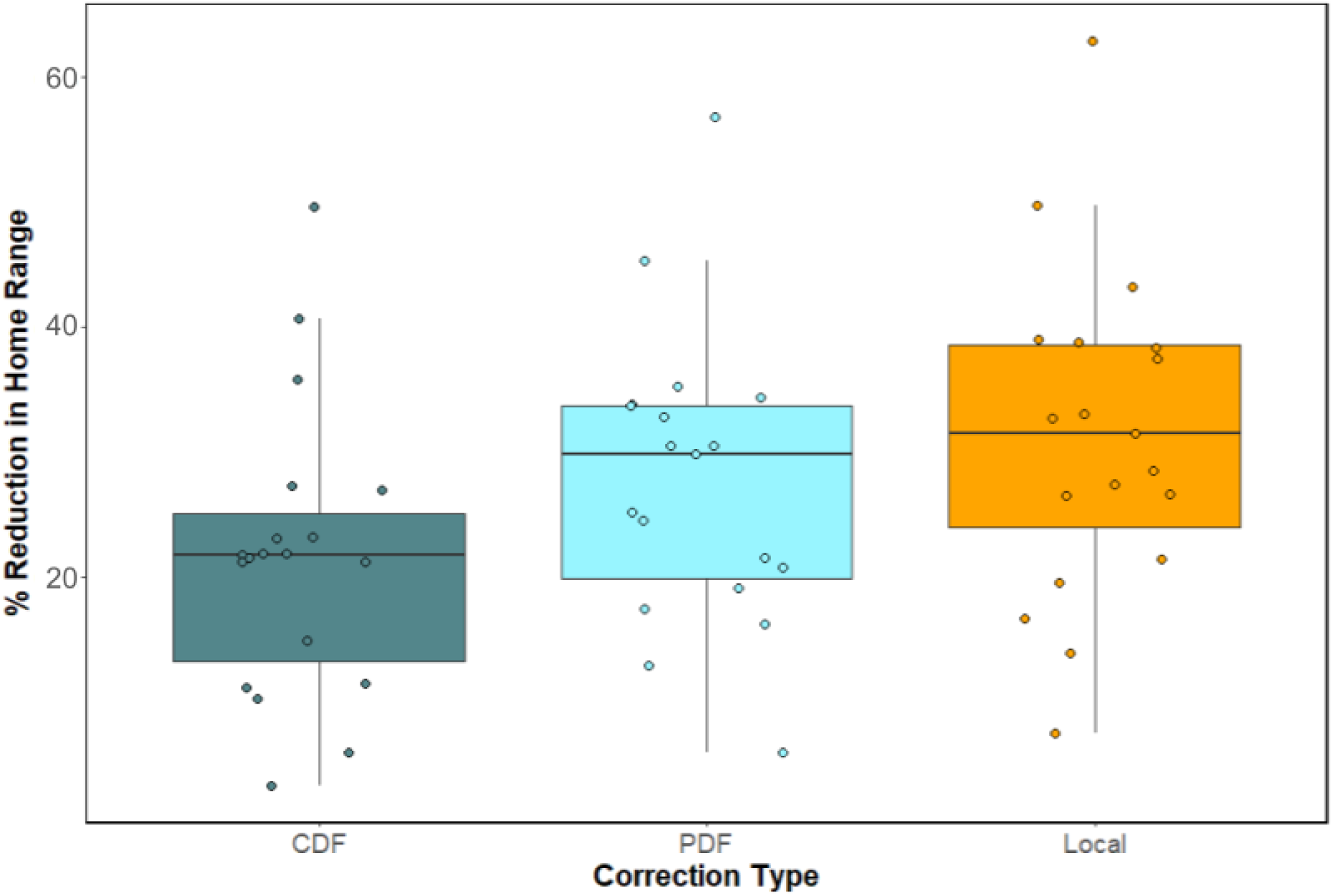
Comparison of the reduction in home range area of lake trout resulting from CDF, PDF and Local corrections. correction constrains redistribution bias, and so reduced the uncorrected home range by the greatest amount in all cases.

**Figure 6:**
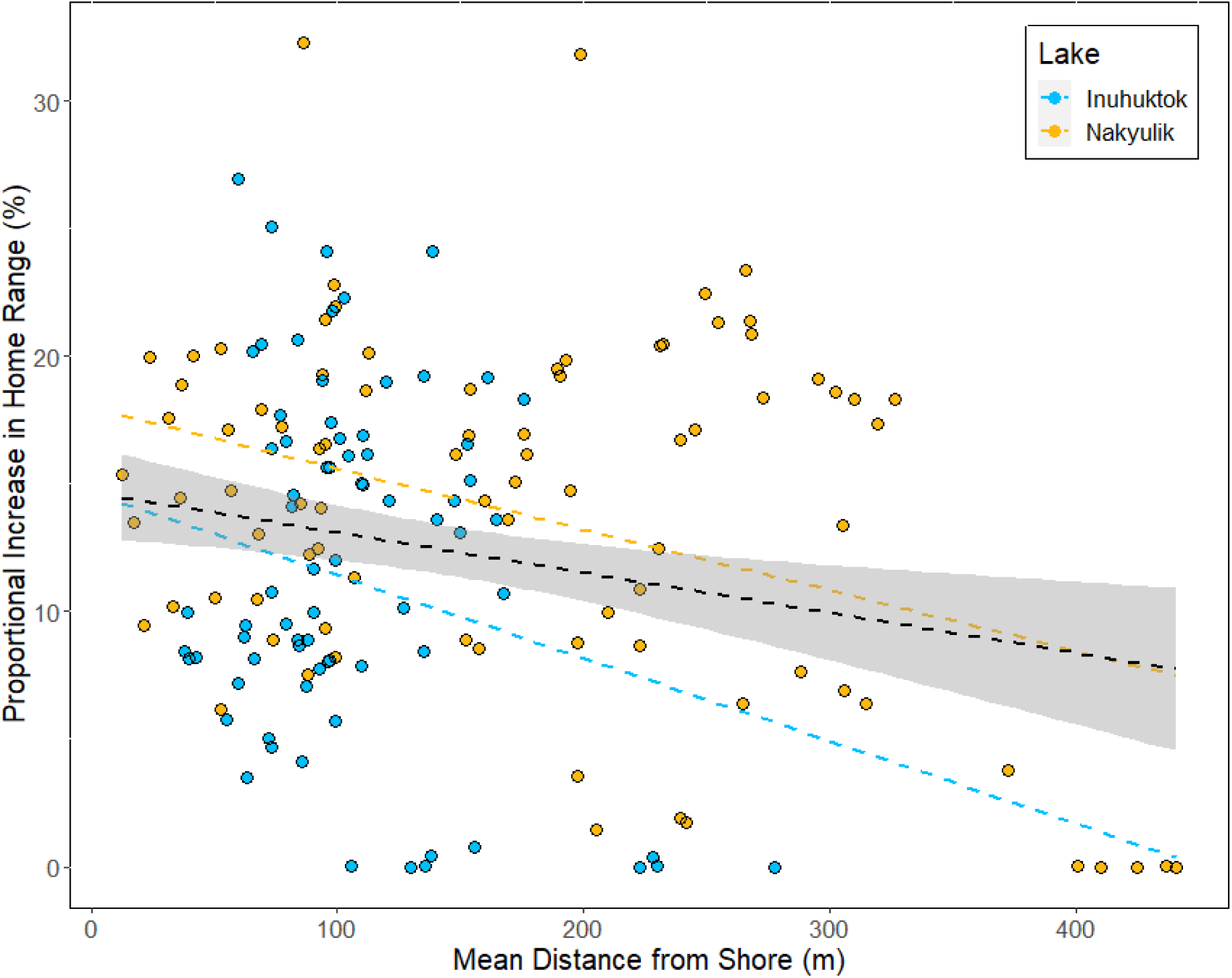
Proportional increase in monthly HR size (comparing post-hoc CDF truncation versus local kernel correction) versus the mean distance fish spent from shore. Colored dashed lines indicate the linear regression for each lake separately,, while the dashed black line indicates the global relationship. Positive bias in CDF corrected home range estimates decreased as fish spent more time further from shore.

**Figure 7:**
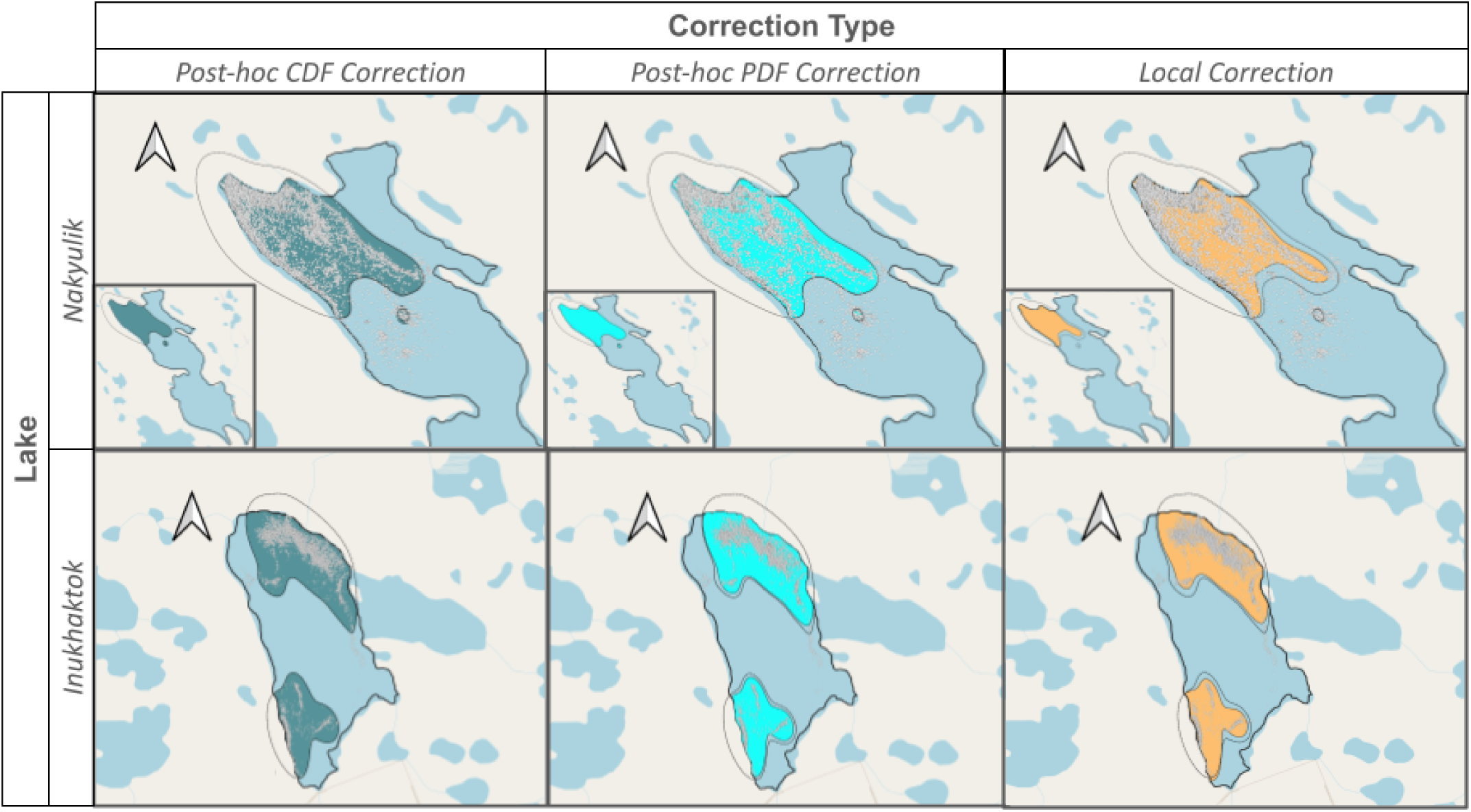
Comparison of lake trout 95% home ranges calculated using data from the full tracking period, from one fish in Nakyulik (top) and Inukhaktok (bottom) lakes, with the boundary enforced via CDF (left), PDF (middle), and local (right) correction. Filled area corresponds to the corrected home range, with dashed lines indicating the size and shape of the uncorrected home range estimate. In these examples, the CDF correction yields a home range which generally adheres to the extent of the uncorrected estimate within the available area, as does the PDF corrected home range in Nakyulik. In Inukhaktok however, the PDF corrected home range retracts behind the uncorrected extent. The locally corrected home range retracts behind the extent of the uncorrected estimate in both lakes.

## Discussion

While the issue of home range spillover has been a point of concern for ecologists for many years, descriptions of the underlying causes and impacts of spillover bias in (A)KDE and the methods used to account for it in home range analysis have been lacking. Here, we first used simulations to demonstrate how local corrections to home range kernels using (A)KDE removed spillover bias and more accurately estimated home range compared to commonly used post-hoc corrections. The redistribution bias of CDF and PDF corrections resulted in higher Bhattacharyya distances between home range estimates and true distributions, while locally corrected home ranges constrained this bias to the area adjacent to the boundary, resulting in more accurate home range distributions in the tracking simulations with both fixed and variable durations. Our real word case study supported that local correction AKDE provided more reliable home range estimates than post hoc corrections where constraint of redistribution bias resulted in significantly smaller trout home ranges.

Avoiding the biases introduced via post-hoc corrections is important when comparing home ranges calculated from individuals whose movement is constrained by different boundaries, or who spend variable amounts of time adjacent to the same boundary (Figs.7,8). For example, the home range of an individual that spends an extended proportion of its time adjacent to an impassable boundary with a large optimal bandwidth will exhibit a large degree of spillover and so yield an erroneously large final home range when CDF corrections are employed. In contrast, a tracked individual exhibiting an identical movement process, but which uses habitat farther from an impassable boundary would have a much smaller CDF-corrected home range. Additionally, where home ranges are used as indicators of periodic changes in space use (e.g. Monk et al. 2023), movements towards and away from impassable boundaries could also result in unpredictable patterns of expansion and contraction in CDF-corrected home ranges, without commensurate changes in scale of space use exhibited by the tracked animal. Finally, where animal movement is wholly constrained by a movement boundary (e.g., an aquatic animal living inside a lake), CDF corrections to home ranges may constrain variation in the area of home ranges present in tracked animals. For example, Fig.8 illustrates how, where sufficient spillover bias is present, CDF corrections can cause home ranges to encompass more of the lake than is used by the individual, potentially masking home range variation between individuals and promoting misinterpretation of habitat-resource use.

**Figure 8:**
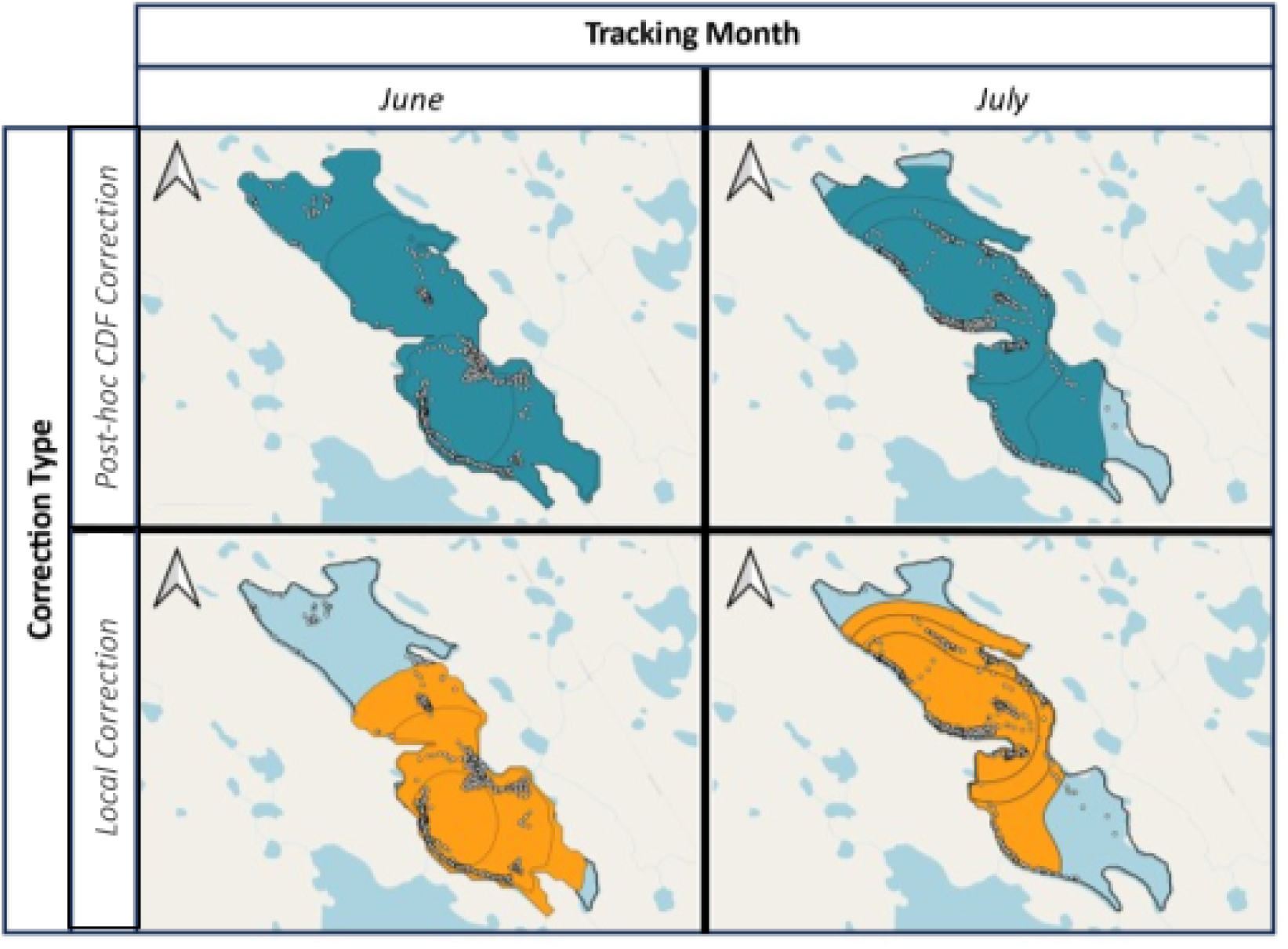
Comparison of monthly HR sizes between June and July calculated for an individual trout within Nakyulik Lake, with boundary enforcement achieved via post-hoc CDF truncation (top) and local kernel truncation (bottom). When using CDF truncation, the large amount of spillover that is redistributed within the available area extends the home range estimate far into the northern lake basin, despite the fish spending little time there in June, and vice-versa in July. In contrast, when using local truncation, the home range is not extended, and more closely adheres to the basin where trout spend more time each month.

*Post-hoc* PDF corrections also introduce positive bias into cell probability densities throughout a home range estimate (Fig 1). Consequently, PDF-corrected home range area estimates will have less bias in home range size compared to those that have had spillover removed via CDF corrections, but will still exhibit positive bias in cell densities throughout the remaining home range extent. PDF corrected home ranges that removed a lot of spillover will therefore yield positively biassed measures of habitat use intensity compared to a home range estimate that exhibited little spillover. In contrast, local corrections to kernels constrain the accumulation of spilled-over probability mass to the home range regions closest to the movement boundary, irrespective of the amount of spillover removed. Local corrections to home range kernels therefore facilitate more direct comparisons of home ranges estimated in environments with different amounts of boundary complexity, and between individuals differently impacted by impassable boundaries.

While spillover bias is impacted by the amount of time an individual spends along an impassable boundary, our variable sample size simulations demonstrate how spillover is particularly exacerbated by low effective sample sizes caused by short tracking durations. As tracking duration increases, spillover decreases as the uncorrected AKDE converges on the true home range, increasing similarity between corrected and uncorrected home range estimates. Consequently, when animals are tracked for long periods of time, uncorrected AKDE home ranges provide less biased area estimates than those provided by any of the corrections investigated here (Fig. 6), but will still show a degree of spillover. However, our simulations showed that locally corrected AKDE exhibited the greatest similarity to the corresponding true distribution across all tracking durations, even when uncorrected AKDEs featured the lowest bias in home range area. As such, the use of locally corrected AKDEs for extended tracking durations will be of particular interest to researchers where probability densities of home range estimates, and their distributions across space are the focus (Fig 6).

The mean proportional spillover for trout home ranges in both Nakyulik and Inukhaktok lakes were higher than in fixed duration simulations, with an average of 36.3%, 25.4% and 18.7% of the uncorrected AKDE spilling over the movement boundary in each context, respectively. As shown in our simulations with variable tracking durations, effective sample size is an important determinant in the amount of spillover; however, effective sample sizes of simulated trajectories were lower than those recorded in both Inukhaktok and Nakyulik trout (mean ±sd effective sample sizes of 33.17±10.07, 552.89±247.8 and 164.38±81.44 in each case, respectively). Accordingly, the larger degree of home range spillover observed in these lakes is likely due to a combination of differing movement behaviour and boundary complexity across scenarios. In our simulations, tracked individuals frequently moved perpendicularly to the impassable boundary, regularly generating positions far enough from the boundary that they did not contribute to spillover. In contrast, trout often moved parallel to the shoreline extensively, such that the majority of their observed locations yielded a corresponding constituent kernel overlapping the boundary (e.g., Figs. 7, 8). These extended associations with a coastal movement boundary are common in fish, with movements often closely paralleling shorelines, exacerbating spillover (e.g., Callihan et al. 2015; Moore et al. 2016; Luo et al. 2020; Hollins et al. 2021). In these cases, while the redistribution bias caused by post-hoc CDF corrections will be exacerbated (Figs 7,8), the performance of PDF vs local corrections will become more similar as the proportion of constituent kernels which extend beyond the coastline increases, minimising the impact of redistribution bias when using PDF correction.

While parallel coastal movements are commonly observed in aquatic taxa, animals exhibit a wide range of responses to encountering impassable boundaries (Jones et al. 2022, Xu et al. 2021). Ungulate responses to anthropogenic movement barriers are the subject of extensive study in terrestrial systems (Jones et al. 2022), including the recognition of six distinct ‘barrier behaviour categories’ (Xu et al. 2021). These behavioural categories include several that result in extended space use near the movement barrier, similar to our trout example. Briefly, these include ‘trace’ and ‘back and forth’ which each represent movement parallel to the barrier with different amounts of directional persistence, and ‘trapped’, representing movement constrained by barriers on all sides (Xu et al. 2021). When conducting home range analyses on animals which exhibit these behavioural responses, the constrained redistribution of probability mass of locally corrected (A)KDE will be particularly beneficial. Where animals ‘bounce off’ of barriers to movement, changing bearing away from the boundary upon its encounter, or where behaviour does not change upon boundary encounter, the redistribution of spillover throughout the entire home range extent may be more appropriate.

Local corrections to kernels are a robust means to account for spillover bias in home range estimation, but they remain corrective, and do not address spillover’s underlying cause. Where animals show strong preferences for specific habitat types, constituent kernels may encompass areas which are available, but unused, by the tracked animal (‘discontinuity bias’; Guo et al. 2019; Peron et al. 2019). While the local corrections to home range kernels demonstrated in this paper address spillover bias across impassable boundaries, home range areas encompassing unfavourable habitat may remain. This bias will be exacerbated where tracking durations are short, and movement tightly constrained by environmental characteristics other than impassable boundaries or where the movement boundary includes narrow peninsulas which extend into the available habitat, but only one side of the peninsula is used.

Addressing discontinuity bias requires a means to inform constituent kernels of the habitat preferences of the tracked animal, and the habitat composition encompassed by that kernel. In this way, the shape of constituent kernels can adhere to the patterns of habitat use and selection exhibited by the animal and prevent extrapolation of home range estimates into unsuitable habitat. Examples of these preventative approaches were implemented by Guo et al. (2019), and Péron (2019), however these are currently inaccessible to most ecologists due to their *ad-hoc* and computationally intensive implementation. Home range estimation approaches that leverage environmental preferences of tracked individuals to constrain the animals’ spatial extents represent powerful tools that could be further developed to address discontinuity bias. The local kernel corrections introduced here provide an improved method for removing spillover bias in home range estimation while avoiding the shortcomings of previous corrections. Local corrections to kernels allow more confident comparison of home range estimates from individuals tracked in environments with contrasting boundary complexity, or among individuals that spend different amounts of time adjacent to impassable boundaries, while mitigating the impacts of redistribution bias associated with CDF correction and to a lesser degree PDF correction. Our new local correction methods also work irrespective of the length of boundary segments relative to the bandwidth, unlike previously available local corrections. Locally corrected AKDE will be particularly beneficial at short/intermediate tracking durations where they greatly reduce biases in estimated home range area. In contrast, uncorrected AKDE showed the least area bias where tracking durations were long, and spillover minimal, suggesting that local corrections may underestimate home range sizes at large sample sizes. However, the consistent low BD between locally corrected AKDE and the true density distribution in simulated data suggests local corrections may retain their advantages at extended tracking durations where patterns of habitat use are of specific interest.

## Supporting information

Supplemental file 1 - Non KDE based HR estimation and spillover

Supplemental file 3 - additional model tables

Supplemental file 4 - point containment in bounded AKDE

Bounded movement simuations and corrections

## Acknowledgements and funding

**JPWH**, and **NEH** supported by University of Windsor, **JPWH**, **LNH**, **BKM**, supported by Fisheries and Oceans Canada, **CHF**, **JMC, JMA, WFF,** supported by University of Maryland, **CHF**, **JMC** supported by CASUS;,Helmholtz Centre for Environmental Research, **CHF** supported by Smithsonian Conservation Biology Institute; University of Central Florida, **JSM** supported by Université Laval, IBIS, RAQ, CEN, **LNH** and **JSM** supported by Fisheries and Oceans Canada (Nunavut Implementation Funds), Nunavut Wildlife Management Board, Polar Knowledge Canada, Ocean Tracking Network, **MJN** supported by University of British Columbia, **JMA** supported by University of Arizona

## Notes

### Competing Interest Statement

The authors have declared no competing interest.

### Summary of Updates

This version updates author affiliations and adds accidentally omitted 'acknowledgements' section.

## References

Alston, J.M., Fleming, C.H., Kays, R., Streicher, J.P., Downs, C.T., Ramesh, T., Reineking, B. and Calabrese, J.M., 2023. Mitigating pseudoreplication and bias in resource selection functions with autocorrelation-informed weighting. Methods in Ecology and Evolution, 14(2), pp.643–654.

Alston, J.M., Fleming, C.H., Noonan, M.J., Tucker, M.A., Silva, I., Folta, C., Akre, T.S., Ali, A.H., Belant, J.L., Beyer, D. and Blaum, N., 2022. Clarifying space use concepts in ecology: range vs. occurrence distributions. BioRxiv, pp.2022–09.

Bhattacharyya, A., 1943. On a measure of divergence between two statistical populations defined by their probability distribution. Bulletin of the Calcutta Mathematical Society, 35, pp.99–110.

Benhamou, S. and Cornélis, D., 2010. Incorporating movement behavior and barriers to improve kernel home range space use estimates. The Journal of Wildlife Management, 74(6), pp.1353–1360.

Burgman, M.A. and Fox, J.C., 2003, February. Bias in species range estimates from minimum convex polygons: implications for conservation and options for improved planning. In Animal Conservation Forum (Vol. 6, No. 1, pp. 19–28). Cambridge University Press.

Calabrese, J.M., Fleming, C.H. and Gurarie, E., 2016. ctmm: An R package for analyzing animal relocation data as a continuous-time stochastic process. Methods in Ecology and Evolution, 7(9), pp.1124–1132.

Callihan, J.L., Harris, J.E. and Hightower, J.E., 2015. Coastal migration and homing of Roanoke River striped bass. Marine and Coastal Fisheries, 7(1), pp.301–315.

Fleming, C.H. and Calabrese, J.M., 2017. A new kernel density estimator for accurate home-range and species-range area estimation. Methods in Ecology and Evolution, 8(5), pp.571–579.

Fleming, C.H., Fagan, W.F., Mueller, T., Olson, K.A., Leimgruber, P. and Calabrese, J.M., 2015. Rigorous home range estimation with movement data: a new autocorrelated kernel density estimator. Ecology, 96(5), pp.1182–1188.

Fleming, C.H., Sheldon, D., Fagan, W.F., Leimgruber, P., Mueller, T., Nandintsetseg, D., Noonan, M.J., Olson, K.A., Setyawan, E., Sianipar, A. and Calabrese, J.M., 2018. Correcting for missing and irregular data in home-range estimation. Ecological Applications, 28(4), pp.1003–1010.

Fleming CH, Drescher-Lehman J, Noonan MJ, Akre TSB, Brown DJ, Cochrane MM, Dejid N, DeNicola V, DePerno CS, Dunlop JN+27 more. 2020. A comprehensive framework for handling location error in animal tracking data. Preprint

Fury CA, Harrison PL, 2008. Abundance, site fidelity and range patterns of Indo-Pacific bottlenose dolphins *Tursiops aduncus* in two Australian subtropical estuaries. Mar Freshw Res 59 :1015–1027

Getz, W.M., Fortmann-Roe, S., Cross, P.C., Lyons, A.J., Ryan, S.J. and Wilmers, C.C., 2007. LoCoH: nonparameteric kernel methods for constructing home ranges and utilization distributions. PloS one, 2(2), p.e207.

Gitzen, R.A. and Millspaugh, J.J., 2003. Comparison of least-squares cross-validation bandwidth options for kernel home-range estimation. Wildlife Society Bulletin, pp.823–831.

Guo, J., Du, S., Ma, Z., Huo, H. and Peng, G., 2019. A Model for animal home range estimation based on the active learning method. ISPRS International Journal of Geo-Information, 8(11), p.490.

Hartel, E.F., Noke Durden, W. & O’Corry-Crowe, G. Testing satellite telemetry within narrow ecosystems: nocturnal movements and habitat use of bottlenose dolphins within a convoluted estuarine system. Anim Biotelemetry 8, 13 (2020).

Hawley, K.L., Rosten, C.M., Christensen, G. and Lucas, M.C., 2016. Fine-scale behavioural differences distinguish resource use by ecomorphs in a closed ecosystem. Scientific reports, 6(1), p.24369.

Hollins, J., Pettitt-Wade, H., Gallagher, C.P., Lea, E.V., Loseto, L.L. and Hussey, N.E., 2022. Distinct freshwater migratory pathways in Arctic char (*Salvelinus alpinus*) coincide with separate patterns of marine spatial habitat-use across a large coastal landscape. Canadian Journal of Fisheries and Aquatic Sciences, 79(9), pp.1447–1464.

Jones, Paul F., Andrew F. Jakes, Scott E. Vegter, and Mike S. Verhage. “Is it the road or the fence? Influence of linear anthropogenic features on the movement and distribution of a partially migratory ungulate.” Movement Ecology 10, no. 1 (2022): 37.

Knight, C.M., Kenward, R.E., Gozlan, R.E., Hodder, K.H., Walls, S.S. and Lucas, M.C., 2009. Home-range estimation within complex restricted environments: importance of method selection in detecting seasonal change. Wildlife Research, 36(3), pp.213–224.

Lapointe, N.W.R., Odenkirk, J.S. and Angermeier, P.L., 2013. Seasonal movement, dispersal, and home range of Northern Snakehead *Channa argus* (Actinopterygii, Perciformes) in the Potomac River catchment. Hydrobiologia, 709, pp.73–87.

Lenth R (2024). _emmeans: Estimated Marginal Means, aka Least-Squares Means_. R package version 1.10.3,

Luo, J., Ault, J.S., Ungar, B.T., Smith, S.G., Larkin, M.F., Davidson, T.N., Bryan, D.R., Farmer, N.A., Holt, S.A., Alford, A.S. and Adams, A.J., 2020. Migrations and movements of Atlantic tarpon revealed by two decades of satellite tagging. Fish and Fisheries, 21(2), pp.290–318.

Lyons, A.J., Turner, W.C. and Getz, W.M., 2013. Home range plus: a space-time characterization of movement over real landscapes. Movement Ecology, 1, pp.1–14.

Mate, I., Barrull, J., Ruiz-Olmo, J., Gosàlbez, J. and Salicrú, M., 2016. Spatial organization and intraspecific relationships of the southern water vole (Arvicola sapidus) in a Mediterranean mountain river: what is the role of habitat quality? Mammal Research, 61, pp.255–268.

Matthiopoulos, J., 2003. Model-supervised kernel smoothing for the estimation of spatial usage. Oikos, 102(2), pp.367–377.

Matthiopoulos, J., 2003a. The use of space by animals as a function of accessibility and preference. Ecological Modelling, 159(2-3), pp.239–268.

Moland, E., Olsen, E.M., Andvord, K., Knutsen, J.A. and Stenseth, N.C., 2011. Home range of European lobster (*Homarus gammarus*) in a marine reserve: implications for future reserve design. Canadian Journal of Fisheries and Aquatic Sciences, 68(7), pp.1197–1210.

Monk, C. T., Power, M., Freitas, C., Harrison, P. M., Heupel, M., Kuparinen, A., Moland, E., Simpfendorfer, C., Villegas-Ríos, D., & Olsen, E. M. (2023). Atlantic cod individual spatial behaviour and stable isotope associations in a no-take marine reserve. Journal of Animal Ecology, 92, 2333–2347. 10.1111/1365-2656.14014

Moorcroft, P.R. and Barnett, A., 2008. Mechanistic home range models and resource selection analysis: a reconciliation and unification. Ecology, 89(4), pp.1112–1119.

Noonan, M.J., Tucker, M.A., Fleming, C.H., Akre, T.S., Alberts, S.C., Ali, A.H., Altmann, J., Antunes, P.C., Belant, J.L., Beyer, D. and Blaum, N., 2019. A comprehensive analysis of autocorrelation and bias in home range estimation. Ecological Monographs, 89(2), p.e01344.

Moore, J.S., Harris, L.N., Kessel, S.T., Bernatchez, L., Tallman, R.F. and Fisk, A.T., 2016. Preference for nearshore and estuarine habitats in anadromous Arctic char (*Salvelinus alpinus*) from the Canadian high Arctic (Victoria Island, Nunavut) revealed by acoustic telemetry. Canadian Journal of Fisheries and Aquatic Sciences, 73(9), pp.1434–1445.

Péron, G., 2019. Modified home range kernel density estimators that take environmental interactions into account. Movement ecology, 7, pp.1–8.

Péron, G., Fleming, C.H., Duriez, O., Fluhr, J., Itty, C., Lambertucci, S., Safi, K., Shepard, E.L. and Calabrese, J.M., 2017. The energy landscape predicts flight height and wind turbine collision hazard in three species of large soaring raptor. Journal of Applied Ecology, 54(6), pp.1895–1906.

Pinheiro J, Bates D, R Core Team (2024). nlme: Linear and Nonlinear Mixed Effects Models. R package version 3.1–166

Signer, J. and Fieberg, J.R., 2021. A fresh look at an old concept: home-range estimation in a tidy world. PeerJ, 9, p.e11031.

Silverman, B.W., 1998. Density estimation for statistics and data analysis. Routledge.

Slaght, J.C., Horne, J.S., Surmach, S.G. and Gutiérrez, R.J., 2013. Home range and resource selection by animals constrained by linear habitat features: an example of Blakiston’s fish owl. Journal of Applied Ecology, 50(6), pp.1350–1357.

Tarjan, L.M. and Tinker, M.T., 2016. Permissible home range estimation (PHRE) in restricted habitats: a new algorithm and an evaluation for sea otters. PLoS One, 11(3), p.e0150547.

Wells, R.S., Schwacke, L.H., Rowles, T.K., Balmer, B.C., Zolman, E., Speakman, T., Townsend, F.I., Tumlin, M.C., Barleycorn, A. and Wilkinson, K.A., 2017. Ranging patterns of common bottlenose dolphins Tursiops truncatus in Barataria Bay, Louisiana, following the Deepwater Horizon oil spill. Endangered Species Research, 33, pp.159–180.

Winner, K., Noonan, M.J., Fleming, C.H., Olson, K.A., Mueller, T., Sheldon, D. and Calabrese, J.M., 2018. Statistical inference for home range overlap. Methods in Ecology and Evolution, 9(7), pp.1679–1691.

Worton, B.J., 1989. Kernel methods for estimating the utilization distribution in home-range studies. Ecology, 70(1), pp.164–168.

Wood, S.N., 2011. Fast stable restricted maximum likelihood and marginal likelihood estimation of semiparametric generalized linear models. Journal of the Royal Statistical Society Series B: Statistical Methodology, 73(1), pp.3–36.

Xu, W., Dejid, N., Herrmann, V., Sawyer, H. and Middleton, A.D., 2021. Barrier Behaviour Analysis (BaBA) reveals extensive effects of fencing on wide-ranging ungulates. Journal of Applied Ecology, 58(4), pp.690–698.

