## Supplemental file 1 - Non KDE based HR estimation and spillover for "Home range spillover in habitats with impassable boundaries: Causes, biases, and corrections using autocorrelated kernel density estimation"

**Appendix S1: Home Range Spillover in Habitats with Impassable Boundaries; Causes, biases, and corrections using AKDE; Supplement 1**

In the main text, the causes and impacts of spillover bias in KDE-based home range (HR) estimation are described, but alternative HR estimators also exist, with contrasting relevance of spillover bias in their use. Accordingly, we discuss three different families of HR estimators, and how they deal with hard movement boundaries in this supplement. For sake of brevity, the following sections focus on the capacity for these estimators to account for movement boundaries, and how that capacity may come at the expense of other aspects of HR estimation, as opposed to their performance as home range estimators in their own right, which is covered more extensively elsewhere in the literature (e.g. Noonan et al. 2019).

***1.Geometric Home Range Estimation***

Geometric home range estimators such as the minimum convex polygon (MCP, Burgman and Fox, 2002), or the local convex hull family of estimators (e.g. T-LoCoH, k-LoCoH, Getz et al. 2007; Lyons et al. 2013) provide an alternative to statistical home range estimation and can avoid the issue of spillover bias when specific conditions are met. Geometric home range estimators construct home ranges from minimum convex hulls, defined as the smallest polygon containing a set of animal relocations in which no internal angle exceeds 180°. The simplest of these is the MCP, which provides a home range estimate constructed from a single hull, encompassing the total extent of animal relocations, while LoCoH home range estimates are composed of multiple local hulls constructed from clusters of neighboring animal relocations. Provided that these animal relocations themselves adhere to the presence of a convex boundary to movement, the extent of range distributions constructed from these hulls will also respect that boundary, avoiding the issue of spillover into unavailable habitat. However, home ranges calculated via geometric home range methods will still result in spillover where boundaries are concave, including where these boundaries are encompassed by the animal’s home range. For example, a geometric home range estimate from a fish swimming in a circular lake will conform to the presence of the shoreline well, however in the presence of a more complex boundary, (e.g. where the shore has several narrow peninsulas extending into the lake body), or in the presence of internal boundaries to movement, such as an island in the center of the lake, the geometric home range estimate can be extrapolated across these inaccessible areas.

There are a number of different implementations of LoCoH (e.g a-LoCoH, k-LoCoH, T-LoCoH), each differentiated by how the number of nearest-neighbor animal locations to be clustered and incorporated into local hulls (Getz *et al.* 2007; Lyons et al. 2013; Dougherty et al. 2018) is determined, while MCP always consists of a single hull. While these geometric home range estimators can account for the presence of convex boundaries to animal movement, they are not optimized for autocorrelated animal tracking data, and so result in asymptotic inconsistencies and underestimation of home range within the available area (Noonan et al. 2019). Of these geometric estimators, home ranges estimated via MCP will yield the largest area extent, while also the exhibiting the least capacity to conform to a complex movement boundary (Getz *et al.* 2007; Lyons et al. 2013). As the number of constituent hulls is increased to better accommodate movement boundaries, the area estimate will continue to decrease below the underestimate of the MCP. Evaluation of the performance of k-LoCoH on bounded movement data (Noonan et al. 2019; Guo et al. 2019) has shown that the negative bias in space use within the available portion of a habitat introduced using convex hull techniques can be greater than the positive bias introduced by a KDE which spills over into areas of impossible use. This is true even in ‘worst case’ scenarios, where regions of high space use by a tracked animal fall along a hard boundary, such that spillover is exacerbated (Noonan et al. 2019). In these instances, statistical (A)KDE with post-hoc corrections to home range extent (main text) will provide much more accurate estimates of home range, provided that the location of these boundaries to movement are already known (Noonan et al. 2019). As post hoc corrections to home ranges estimated via KDE provide more accurate home range estimates than geometric methods, geometric home range estimation is most applicable in instances where animal movement data shows low/absent autocorrelation and are bound by impassable convex boundaries of unknown position/shape, and so cannot be accounted for *post-hoc.*

***2. Mechanistic Home Range Analysis (MHRA)***

Conceptually, MHRA treats an animal’s home range as a trait which manifests from the cumulative outcome of a tracked animal’s suite of behaviors, first fitting mechanistic movement models to an animal’s trajectory then scaling these underlying movement processes ‘upwards’ to provide an estimate of space use (Moorcroft and Barnett, 2008; Borger et al. 2008). These underlying movement models can parameterize a range of behaviors, (e.g. responses to environmental and social factors) which ultimately shape an animal’s home range, including anchoring home ranges to focal points such as dens or haul-out sites (Liukkonen et al. 2018; Moorcroft et al, 2006), avoidance of conspecific territories at the edges of home ranges, (Briscoe et al. 2002), and selection/avoidance of specific resources or environmental conditions, including restricting foraging to high density resource patches (Moorcroft and Barnett, 2008), or shaping movement corridors in accordance with surrounding landscape topography (Bateman et al. 2015; Eisaguirre et al. 2022). Analytically, MHRA constructs home range estimates by first characterizing fine-scale movement behaviours as a stochastic movement process, defined by a redistribution kernel which specifies the probability of a tracked animal moving from its current position to the surrounding area within the time interval preceding the next recorded animal position (Moorcroft and Barnett, 2008). In addition to providing a set of movement rules describing the probability of an individual moving from position to position as a function of time and space, additional coefficients can be used to link kernel propagation to specific behaviors or environmental conditions. For example, a reorientation coefficient can direct the propagation of redistribution kernels towards focal points or otherwise direct the spread of the kernel towards preferentially utilized areas as defined by an underlying resource selection function fit to the animal’s relocations. The collective pattern formed from these redistribution kernels can then be described using a probability density function, the extent of which provides an estimate of the tracked animal’s home range.

The capacity for MHRA to generate home range estimates that are informed by patterns of habitat selection has direct implications for avoiding the issue of discontinuity bias, and the spillover it causes. For example, MHRA approaches can include conspecific avoidance responses, biasing propagation of redistribution kernels towards their home range centers after encountering the boundaries of conspecific territories (Moorcroft and Lewis, 2006). Provided that tracked animal relocations sufficiently capture these patterns of habitat selection, and that relevant underlying environmental data (e.g., elevation or bathymetry) are available to the subsequent MHRA, then it is possible to derive probability density functions for redistribution kernels which adhere to the ‘movement rule’ of never crossing a hard boundary, or redirecting movement away from a repelling boundary such as a busy road. The resulting home range estimate will consequently also adhere to the presence of boundaries. However, while such an approach is intuitive, topographical and/or bathymetric data have yet to be used to specifically constrain home range estimates to within a given boundary to movement within an MHRA framework.

***3. Lattice-based diffusion models***

Lattice-based home range estimation was developed by Barry and McIntyre (2011) as an alternative to KDE for use where tracked animals move within irregular boundaries. When using lattice-based home range estimation, animal relocations are discretised to adhere to a node within a regularly spaced lattice of positions with a resolution defined by the user (Barry and McIntyre, 2011). These nodes are bidirectionally connected to one another via user defined vertical, horizonal, and diagonal links, such that nodes can be directly connected to a maximum of eight adjacent positions. Where an impassable boundary to movement intersects these directional links, the link is removed, so that nodes on either side of impassable boundaries do not join. The distribution of rediscretised animal positions across this lattice is then used to assign an initial probability mass to each node, such that nodes where more positions were recorded receive higher probability, and the probability across all nodes sums to one. This initial probability density is then propagated outwards towards adjacent nodes along the user defined lattice-network via random walks, with probabilities of transitions between nodes informed by node transition matrix, generated using a user defined ‘M’ term, governing the probability of the random walk remaining in the same location after one step. The degree to which the initial probability mass is spread to adjacent nodes is controlled via a ‘k’ term, with higher values corresponding to a greater number of steps within the random walk, and thus a smoother probability density which extends further from the initial set of observations. In this way, the ‘k’ parameter in lattice-based home range estimation can be considered analogous to the bandwidth in KDE. As the random walks implemented in lattice-based home range estimation can only move along user defined links between nodes, the shape of the resulting PDF after ‘k’ steps will adhere to the presence of boundaries intersecting nodes which would have otherwise been adjacent.

The value of ‘k’ used in lattice-based home range estimation will be a determinant factor in the size and shape of the resulting PDF, similarly to that of bandwidth used in KDE, and is therefore optimized analytically (Barry and McIntyre, 2011). K-optimization by Barry and McIntyre(2011) was conducted using leave-one-out cross validation (LOOCV), and as such assumes the underlying node-transitions are IID, potentially leading to underestimation of ‘k’ when performed on more autocorrelated animal tracking data, as will generally be the case (Noonan et al. 2019). In addition to the method of k-optimisation assuming IID node-transitions the diffusive propagation of probability mass controlled via random walks assumes that the movement of the tracked animal between nodes is Brownian, i.e., that animal velocities are uncorrelated, and space use is otherwise unbounded (Einstein, 1905; Horne et al. 2007; Calabrese et al. 2016). While uncorrelated velocities in animal movement is often observed in telemetry datasets, the assumption of unbound movement is incongruous with the concept of home range (Noonan et al. 2019), and transitions between nodes would be more appropriately represented by an Ornstein Uhlenbeck process, which also exhibits uncorrelated movement velocities, but with a central tendency, facilitating more accurate HR estimation.

**References**

Barry, R.P. and McIntyre, J., 2011. Estimating animal densities and home range in regions with irregular boundaries and holes: A lattice-based alternative to the kernel density estimator. Ecological Modelling, 222(10), pp.1666-1672.

Bateman, A.W., Lewis, M.A., Gall, G., Manser, M.B. and Clutton‐Brock, T.H., 2015. Territoriality and home‐range dynamics in meerkats, Suricata suricatta: a mechanistic modelling approach. Journal of Animal Ecology, 84(1), pp.260-271.

Börger, L., Dalziel, B.D. and Fryxell, J.M., 2008. Are there general mechanisms of animal home range behaviour? A review and prospects for future research. Ecology letters, 11(6), pp.637-650.

Briscoe, B.K., Lewis, M.A. and Parrish, S.E., 2002. Home range formation in wolves due to scent marking. Bulletin of Mathematical Biology, 64(2), pp.261-284.

Burgman, M.A. and Fox, J.C., 2003, Bias in species range estimates from minimum convex polygons: implications for conservation and options for improved planning. In Animal Conservation Forum (Vol. 6, No. 1, pp. 19-28). Cambridge University Press.

Dougherty, E.R., de Valpine, P., Carlson, C.J., Blackburn, J.K. and Getz, W.M., 2018. Commentary to: A cross-validation-based approach for delimiting reliable home range estimates. Movement ecology, 6(1), pp.1-5

Eisaguirre, J.M., Booms, T.L., Barger, C.P., Lewis, S.B. and Breed, G.A., 2022. Demographic partitioning of dynamic energy subsidies revealed with an Ornstein–Uhlenbeck space use model. Ecological Applications, 32(4), p.e2542.

Getz, W.M., Fortmann-Roe, S., Cross, P.C., Lyons, A.J., Ryan, S.J. and Wilmers, C.C., 2007. LoCoH: nonparameteric kernel methods for constructing home ranges and utilization distributions. PloS one, 2(2), p.e207.

Guo, J., Du, S., Ma, Z., Huo, H. and Peng, G., 2019. A Model for animal home range estimation based on the active learning method. ISPRS International Journal of Geo-Information, 8(11), p.490.

Liukkonen, L., Ayllón, D., Kunnasranta, M., Niemi, M., Nabe-Nielsen, J., Grimm, V. and Nyman, A.M., 2018. Modelling movements of Saimaa ringed seals using an individual-based approach. Ecological Modelling, 368, pp.321-335.

Lyons, A.J., Turner, W.C. and Getz, W.M., 2013. Home range plus: a space-time characterization of movement over real landscapes. Movement Ecology, 1, pp.1-14.

Moorcroft, P.R., Lewis, M.A. and Crabtree, R.L., 2006. Mechanistic home range models capture spatial patterns and dynamics of coyote territories in Yellowstone. Proceedings of the Royal Society B: Biological Sciences, 273(1594), pp.1651-1659.

Moorcroft, P.R. and Barnett, A., 2008. Mechanistic home range models and resource selection analysis: a reconciliation and unification. Ecology, 89(4), pp.1112-1119.

Noonan, M.J., Tucker, M.A., Fleming, C.H., Akre, T.S., Alberts, S.C., Ali, A.H., Altmann, J., Antunes, P.C., Belant, J.L., Beyer, D. and Blaum, N., 2019. A comprehensive analysis of autocorrelation and bias in home range estimation. Ecological Monographs, 89(2), p.e01344.
