## Supplemental file 3 - additional model tables for "Home range spillover in habitats with impassable boundaries: Causes, biases, and corrections using autocorrelated kernel density estimation"

| **Model** | **Term** | **Estimate** | **SE** | **t** | **p** | ***r_m_^2^*** | ***r_c_^2^*** |
| --- | --- | --- | --- | --- | --- | --- | --- |
| Proportional overestimation of CDF vs Locally corrected HR~Mean distance from shore (Simulation) | Intercept | 1.068 | <0.001 | 4444.23 | <0.001 | NA | NA |
|  | Distance | -0.001 | <0.001 | -15.04 | <0.001 |  |  |
| Reduction in HR size~Correction Type + (1\|ID) | Intercept | 0.217114 | 0.027834 | 7.8 | <0.001 | 0.098 | 0.97 |
|  | CDF vs Local | 0.095958 | 0.007168 | 8.36 | <0.001 |  |  |
|  | PDF vs CDF | 0.059914 | 0.007168 | 13.39 | <0.001 |  |  |
|  | CDF vs PDF (post hoc test) | -0.0599 | 0.00717 | -8.359 | <0.001 | NA | NA |
|  | CDF vs Local (post hoc test) | -0.096 | 0.00717 | -13.387 | <0.001 | NA | NA |
|  | PDF vs Local (post hoc test) | -0.036 | 0.00717 | -5.029 | <0.001 | NA | NA |
| Proportional overestimation of CDF vs Locally corrected HR~Mean distance from shore (Nakyulik) | Intercept | 117.971 | 1.3 | 91.081 | <0.001 | NA | NA |
|  | Distance | -0.023 | 0.006 | -3.967 | <0.001 |  |  |
| Proportional overestimation of CDF vs Locally corrected HR~Mean distance from shore (Inuhuktok) | Intercept | 114.658 | 1.623 | 70.65 | <0.001 | NA | NA |
|  | Distance | -0.037 | 0.019 | -2.355 | 0.022 |  |  |

**Table S3.1;**Results of linear and linear mixed effects models examining the relationship between distance from shore and proportional reductions in HR size caused by local truncation, and differences in HR size between different truncation techniques.

| **Model** | **Term** | **Estimate** | **SE** | ***t*** | ***p*** | ***k*** |
| --- | --- | --- | --- | --- | --- | --- |
| % Accuracy~Tracking duration + Correction Type | Intercept | 10.387 | 0.1241 | 83.69 | <0.001 | 12 |
|  | CDF vs PDF | -2.932 | 0.1755 | -16.71 | <0.001 |  |
|  | CDF vs Local | -5.899 | 0.1755 | -33.61 | <0.001 |  |
| Smooth | **Term** | Eff.df | Ref.df | ***F*** | ***p*** |  |
|  | Tracking Dur. | 9.998 | 10 | 9086 | <0.001 |  |
| **Model** | **Term** | **Estimate** | **SE** | ***t*** | ***p*** | ***k*** |
| Bhattacharyya Distance~Tracking duration + Correction Type | Intercept | 0.165 | 0.002 | 81.723 | <0.001 | 12 |
|  | CDF vs PDF | 0.001 | 0.003 | 0.197 | <0.001 |  |
|  | CDF vs Local | -0.3 | 0.003 | -10.453 | <0.001 |  |
| Smooth | **Term** | Eff.df | Ref.df | ***F*** | ***p*** |  |
|  | Tracking Dur. | 9.923 | 9.993 | 509.9 | <0.001 |  |

**Table S3.2;**Results of generalised additive models examining the relationship between % accuracy of HR estimates and Bhattacarrya distance between HR estimates and the true distribution forHR corrections and tracking durations,
