## Supplemental file 4 - point containment in bounded AKDE for "Home range spillover in habitats with impassable boundaries: Causes, biases, and corrections using autocorrelated kernel density estimation"

**Appendix S4: Point Containment in Bounded AKDE**

In the main text of this paper, we demonstrate the performance of spillover removal via post-hoc CDF, PDF, and newly introduced local corrections using simulations, and go on to demonstrate the impacts of these different corrections on a real world example. In our simulations, both true area and density distributions were known, allowing estimation of the bias of the different corrections, but this was not possible for our case study on lake trout. In this supplement, we demonstrate differences in point-containment of trout home ranges to provide additional context for the strengths and weaknesses of the different corrections. An unbiased 95% home range area estimate should contain 95% of animal positions observed over the tracking period, with area estimates containing >95% or <95% indicative of area overestimates and underestimates, respectively. When impassable movement barriers are not present, and so boundary corrections are unnecessary, the final shape and size of the home range estimate is determined by the bandwidth, in turn determining the proportion of animal positions contained by the home range estimate. However, when correcting for home range spillover, removed home range extent is redistributed within the available home range area (redistribution bias), in turn impacting the final size and shape of home range. As post-hoc CDF, PDF and local kernel corrections each redistribute removed probability mass differently, they may also introduce bias into the remaining home range area, impacting the proportion of positions contained within the estimate.

To investigate the impact of post-hoc and local corrections on point containment of corrected home range estimates, we determined the proportion of recorded trout locations contained within corresponding uncorrected home ranges estimated via AKDE using positions from the full study period.

We then enforced the presence of an impassable boundary using post-hoc and local corrections. The resulting deviation from the target 95% point containment was then used as an indicator of the bias introduced by the boundary corrections, and compared using ANOVA.

Mean bias in point containment showed no significant difference between the different correction approaches (F_(3,76)_=0.58, p=0.63), although mean bias for uncorrected and CDF and PDF corrected estimates was positive (each containing an average of 96.1±0.16 and 96.5±0.16 and 95.2% ±0.16, respectively), but was negative for local corrections (93.9% ± 0.24). In terms of proportional point inclusion, post-hoc PDF correction therefore showed the least amount of mean bias (0.2%), followed by local (1.1%) and CDF (1.5%) corrections.

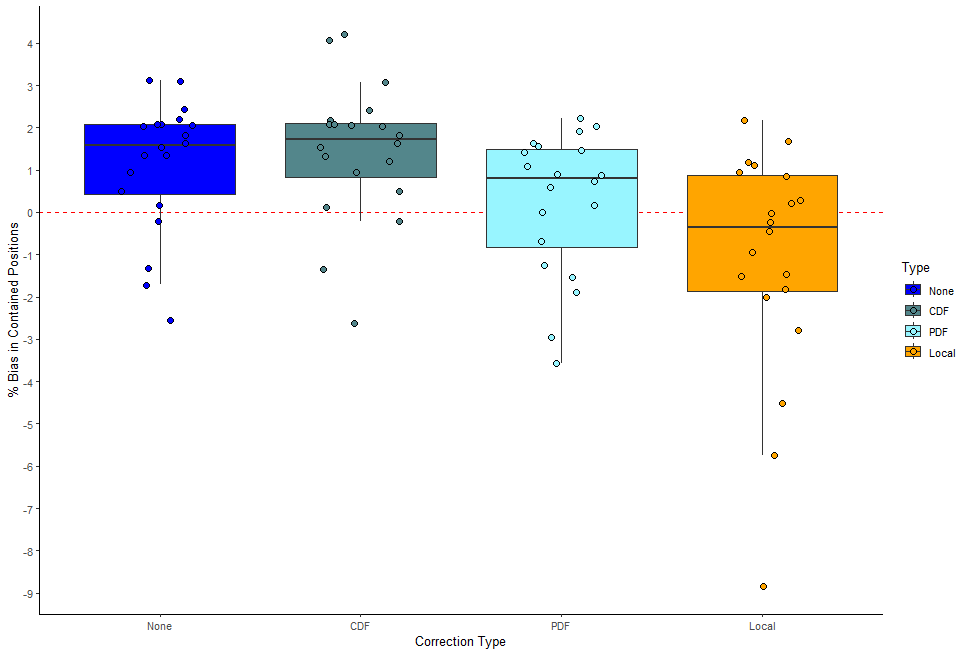

Figure S4.1; *Comparison of the deviation from 95% point containment in uncorrected, CDF and PDF post-hoc corrected, and locally corrected home ranges estimated for lake trout. The dashed red line corresponds to 95% point containment, or 0% bias. Compared to the uncorrected home range, CDF corrected home range estimates generally contained a similar number of points, both exhibiting a slight positive bias in proportion of points contained. Redistribution bias resulting from CDF correction can increase the proportion of contained positions compared to the uncorrected home range. Post-hoc PDF correction can result in a contracted home range within the available area compared to the uncorrected home range, resulting in fewer positions contained within the estimate. Local correction results in the largest reduction in home range area, and so resulted in the least number of positions contained within the home range extent, and also showed the greatest deviance from 95% point containment of the correction approaches. Despite this, there was no significant difference in % point containment bias between the different correction approaches.*
