## Supplementary material for "Home range spillover in habitats with impassable boundaries: Causes, biases, and corrections using autocorrelated kernel density estimation": Bounded movement simuations and corrections

##############################################################
#Simulation of movement constrained by an impassable boundary#
##############################################################
#Follows Noonan et al. 2019 

library(ctmm)
library(sp)
library(terra)

#Define the sample schedule 
#1 day in seconds (also used to define home range crossing time)
ds<-86400
#Tracking duration (days)
nd<-32
#Number of samples per day 
pd<-8
#sampling time (in seconds)
st<-1:(nd*pd)*(ds/pd)

#Specify a base OU model 
MODEL.OU<-ctmm(mu=c(0,0),isotropic=TRUE,sigma=1,tau=c(ds), profile =T)
MODEL.OU@info$projection<-'+proj=utm +zone=1' 
#Define the unbounded model for x-axis positions
FREE.OU.X<-ctmm(mu=MODEL.OU$mu, sigma=MODEL.OU$sigma, tau=MODEL.OU$tau)#create model using parameters of 'MODEL.OU' to generate x positions
#Define the bounded model for the y-axis positions 
Bounded.Y.OU=ctmm(mu=MODEL.OU$mu, sigma=sqrt(MODEL.OU$sigma/4), tau = 2*MODEL.OU$tau)#create model using parameters of 'MODEL.OU' to generate bound 'y' positions
#give these the same projection, required for the local corrections in AKDE 
#FREE.OU.X@info$projection<-'+proj=utm +zone=1' 
#Bounded.Y.OU@info$projection<-'+proj=utm +zone=1' 

#Simulate data from the unbounded and bounded and unbounded models 
Bounded.OU<-ctmm:::pseudonymize(data.frame(t = st, 
                                           x = simulate(FREE.OU.X, t = st)$x, 
                                           y = (simulate(Bounded.Y.OU, t = st)$y^2 + simulate(Bounded.Y.OU, t = st)$y^2)), 
                                proj = projection(MODEL.OU@info$projection))

#Convert to telemetry object
Bounded.OU<-ctmm:::new.telemetry(Bounded.OU[[1]])
uere(Bounded.OU)<-NULL

#Fit OU movement model to bounded tracjectory 
Bounded.OU_fit<-ctmm.fit(Bounded.OU, MODEL.OU)

#Estimate uncorrected home range via AKDE
Whole_AKDE <- akde(Bounded.OU, Bounded.OU_fit)

#create boundary shape for corrections 
coords_1 = matrix(c(-20,0,
                    -20,20,
                    20,20,
                    20,0), 
                  ncol = 2, byrow = TRUE)

bound = Polygon(coords_1)
bound = SpatialPolygons(list(Polygons(list(bound), ID = "a")))
crs(bound)<-'EPSG:32601'#create boundary with same projection
#check home range, boundary, and positions 
plot(Whole_AKDE)
plot(Bounded.OU, add=T, error=F)
plot(bound, add=T)

################################
#####Locally corrected AKDE#####
################################
Local_AKDE<-akde(Bounded.OU,Bounded.OU_fit,weights=F, debias=F, smooth=F, trace=3, fast=T, PC='direct', SP=bound, SP.in = TRUE) 
plot(Local_AKDE)

################################
#####Post-hoc CDF correction####
################################
library(sf)
bound <- st_as_sf(bound)
CDF_C_AKDE <- Whole_AKDE
#function for CDF corrections
update_CDF <- function(CDF_C_AKDE, bound) {
  # Create a grid of points corresponding to CDF matrix
  x_vals <- seq(min(CDF_C_AKDE$r$x), max(CDF_C_AKDE$r$x), length.out = ncol(CDF_C_AKDE$CDF))
  y_vals <- seq(min(CDF_C_AKDE$r$y), max(CDF_C_AKDE$r$y), length.out = nrow(CDF_C_AKDE$CDF))
  points_df <- expand.grid(x = x_vals, y = y_vals)
  # Convert the grid of points to a spatial object
  points_sf <- st_as_sf(points_df, coords = c("x", "y"), crs = st_crs(bound))
  # check which CDF cells fall within boundary 
  contained <- st_contains(bound, points_sf, sparse = FALSE)
  # Extract the cells outside the boundary
  not_contained_indices <- which(!contained)
  # Initialize the CDF matrix
  updated_CDF <- CDF_C_AKDE$CDF
  # Update the CDF matrix based on containment result
  if (length(not_contained_indices) > 0) {
    updated_CDF[not_contained_indices] <- 1
  }
  return(updated_CDF)
}
#correct CDF to Boundary 
CDF_C_AKDE$CDF <- update_CDF(CDF_C_AKDE, bound)

#Calculate corresponding PDF from corrected CDF 
CDF <- CDF_C_AKDE$CDF
dr <- CDF_C_AKDE$dr
DIM <- dim(CDF)
CDF <- c(CDF)
CDF <- sort(CDF, index.return = TRUE)
IND <- CDF$ix
CDF <- CDF$x
PMF <- ctmm:::cdf2pmf(CDF)
PMF[IND] <- PMF
PMF <- array(PMF, DIM)
PDF <- PMF / prod(dr)
CDF_C_AKDE$PDF <- PDF

#corrected CDF
plot(CDF_C_AKDE, DF='CDF')
#corresponding PDF
plot(CDF_C_AKDE, DF='PDF')

################################
#####Post-hoc PDF correction####
################################
PDF_C_AKDE <- Whole_AKDE
#function for PDF correction
update_PDF <- function(PDF_C_AKDE, bound) {
  # Create a grid of points corresponding to PDF matrix
  x_vals <- seq(min(PDF_C_AKDE$r$x), max(PDF_C_AKDE$r$x), length.out = ncol(PDF_C_AKDE$PDF))
  y_vals <- seq(min(PDF_C_AKDE$r$y), max(PDF_C_AKDE$r$y), length.out = nrow(PDF_C_AKDE$PDF))
  points_df <- expand.grid(x = x_vals, y = y_vals)
  # Convert the grid of points to a spatial object
  points_sf <- st_as_sf(points_df, coords = c("x", "y"), crs = st_crs(bound))
  # check which PDF cells fall within boundary 
  contained <- st_contains(bound, points_sf, sparse = FALSE)
  # Extract the indices of points that are not contained
  not_contained_indices <- which(!contained)
  # Initialize the PDF matrix
  updated_PDF <- PDF_C_AKDE$PDF
  # Update the PDF matrix based on containment result
  updated_PDF[not_contained_indices] <- 0
  return(updated_PDF)
}

PDF_C_AKDE$PDF <- update_PDF(PDF_C_AKDE, bound)
#renormalise and calculate CDF
PDF_C_AKDE$PDF=PDF_C_AKDE$PDF/(sum(PDF_C_AKDE$PDF)*prod(PDF_C_AKDE[["dr"]]))
PDF_C_AKDE$PMF=PDF_C_AKDE$PDF*prod(PDF_C_AKDE[["dr"]])#you can store the PMF within the UD object
PDF_C_AKDE$CDF=ctmm:::debias.volume(PDF_C_AKDE$PMF)$CDF

plot(PDF_C_AKDE, DF='PDF')
plot(PDF_C_AKDE, DF='CDF')
